# Tuning the open-close equilibrium of Cereblon with small molecules influences protein degradation

**DOI:** 10.64898/2025.12.19.695617

**Authors:** Suzanne O’Connor, Zoe J. Rutter, Angus D. Cowan, Markus Zeeb, Florian Binder, Yuting Cao, Sohini Chakraborti, Stefan Djukic, Leonhard Geist, Elizabeth Hogg, Giorgia Kidd, Matthias Krumb, Simon Langer, Denis Schmidt, Liam Martin, Elisha H. McCrory, Giacomo Padroni, Ilaria Puoti, Luke Simpson, Manon Sturbaut, Lisa Crozier, Vesna Vetma, Qingzhi Zhang, Kirsten McAulay, Theodor Theis, Alessio Ciulli

## Abstract

Most PROTACs and molecular glue degraders currently approved or in clinical trials recruit Cereblon (CRBN) as the ubiquitin E3 ligase. Upon binding ligands and molecular glues, CRBN undergoes a significant structural rearrangement from an open to closed state, defined by the positioning of the thalidomide-binding domain (TBD) with respect to the Lon domain. However, the exact molecular basis for this ligand-induced conformational change and its implication to neo-substrate degradation remain elusive. During our campaign to discover novel CRBN binders, we found hits exhibiting distinct biophysical behaviour from classical thalidomide-based ligands. By combining orthogonal biophysical methods of differential scanning fluorimetry, isothermal titration calorimetry, and small-angle X-ray scattering, supported by X-ray crystallography and cryo-EM structures of ligand-bound complexes, we classify CRBN binders between those that can induce CRBN closure, and those that cannot. Mutational studies identify key residues in the CRBN ligand binding pocket and N-terminal belt that are essential for the Lon and TBD domains to come together in the closed state. Finally, we show that the probability to yield active degrader molecules is greatly influenced by whether binders can or cannot induce CRBN closure. Together, our study reveals new molecular insights into the structural basis for how CRBN open-closed equilibrium is directly modulated by compound binding and impact target degradability by CRBN, with important implications to the design of PROTACs and molecular glue degraders.

## Introduction

The ubiquitin-proteasome system (UPS) tags proteins for proteasomal degradation through polyubiquitination, which is initiated by attachment of ubiquitin to amino acid residues, typically lysine, on the protein surface.^1^ This process is facilitated by E3 ubiquitin ligases which interact with and position the protein substrates for ubiquitination. A growing variety of modalities, including proteolysis targeting chimeras (PROTACs) and molecular glue degraders (MGDs), hijack this natural system to tag proteins of interest with ubiquitin for targeted protein degradation (TPD). PROTACs are heterobivalent molecules which consist of an E3 ligase ligand connected to a ligand for a target protein via a linker, whilst molecular glues are typically monovalent. These molecules induce formation of a ternary complex between the E3 ligase, PROTAC or MGD, and target protein, leading to ubiquitination and subsequent degradation of the protein by the proteosome. This is a powerful mechanism of action that has expanded the druggable proteome. The Cullin-4 RING (CRL4) ligase cereblon (CRBN) complex, plays a critical role in marking proteins for degradation via the proteasome and has been widely exploited in TPD. In a natural setting, substrate specificity is mediated by the substrate receptor subunit CRBN, which is a component of the CRL4 complex (consisting of the Cullin-4 scaffold, the DNA damage-binding protein 1 (DDB1) adaptor protein, and RING finger protein (ROC1)). CRBN consists of three main folded domains: an N-terminal Lon protease-like domain (Lon), a central helical bundle (HB), and a C-terminal thalidomide-binding domain (TBD). CRBN adopts either an ‘open’ or ‘closed’ conformation that is influenced by ligand binding. In its apo form, CRBN exclusively adopts the open conformation, where the Lon domain and TBD do not interact, and the TBD sensor loop and N-terminal belt upstream of the Lon domain are disordered. Ligand binding to the TBD induces a conformational change to the closed conformation, where the TBD and Lon domains pack tightly against each other through ordering of the sensor loop and N-terminal belt at a large newly formed interface (Figure 1).

**Figure 1.**
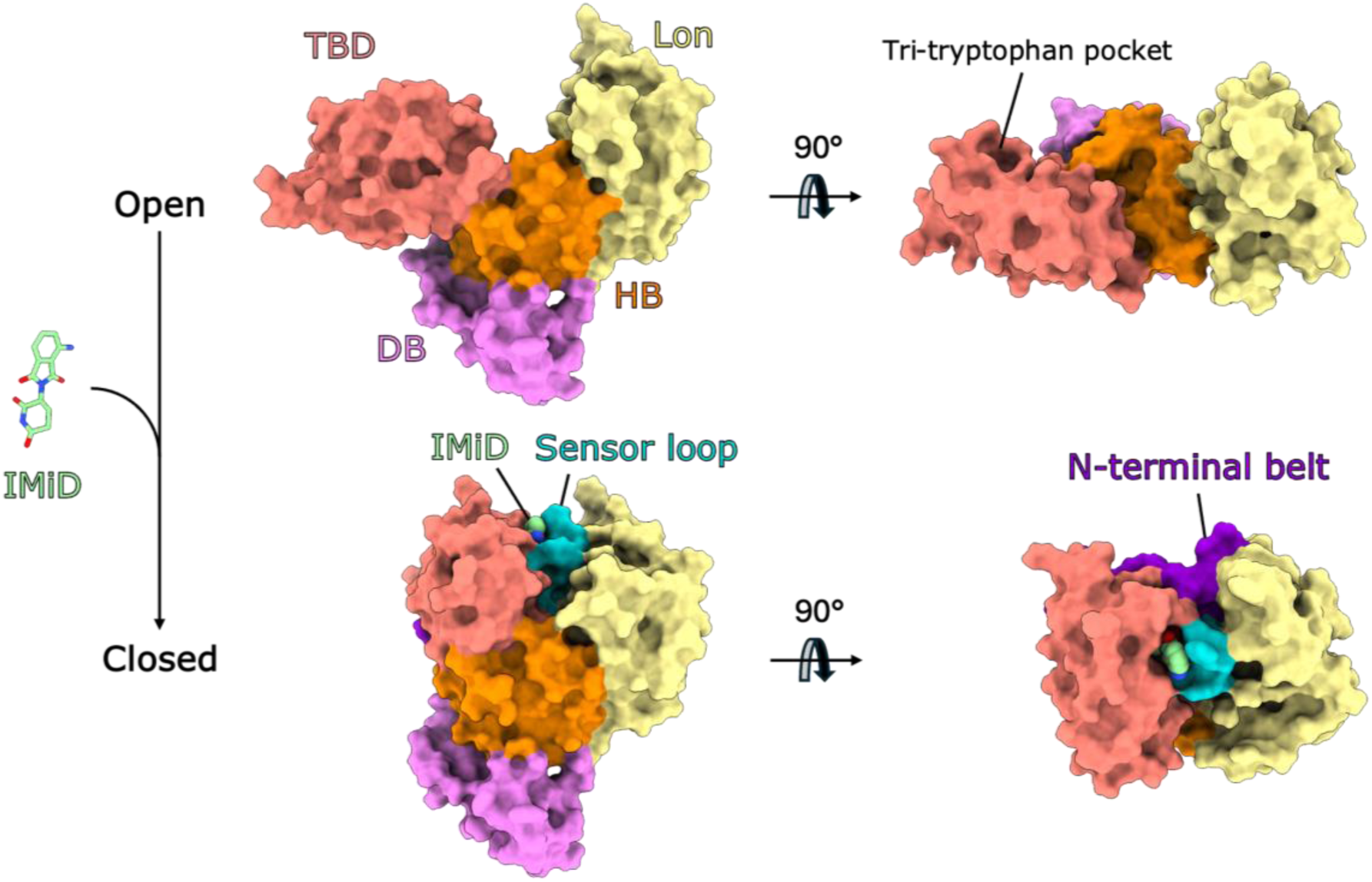
Ligand-induced structural transition of CRBN from the open to closed conformation. Open state (top panels, PDB code 8CVP) and closed states (bottom panels, PDB code 8D81) are shown in surface representation. Binding of ligands such as the IMiD pomalidomide (green) to the tri-tryptophan pocket of the TBD (salmon) induces reorientation of the TBD, ordering of the sensor loop (teal) and N-terminal belt (purple), such that the bound IMiD and sensor loop sit enclosed by the Lon domain (yellow), TBD and N-terminal belt. The DDB1-binding (DB) region and helical bundle (HB) are coloured violet and orange, respectively.

IMiDs (immunomodulatory imide drugs) were the first class of CRBN binders discovered (Figure 1) and are structural mimics of CRBN’s natural degron, the C-terminal cyclic imides that were identified subsequently ^2,3,4^ IMiDs such as thalidomide, lenalidomide and pomalidomide bind with high affinity to the tri-tryptophan cage pocket within the TBD, which is minimally sufficient for ligand binding. Binding at this site also induces an elaborate structural transition in CRBN to the closed conformation, which has been implicated in its function (Figure 1).^5^ Upon ligand binding, the disordered TBD sensor loop, previously engaged with beta-propeller A of DDB1, folds into a beta-hairpin that sits adjacent to the ligand bound to the tri-tryptophan pocket. The ligand-bound TBD-sensor loop module undergoes a ∼25 Å shift and ∼80° rotation such that the sensor loop and bound IMiD are enclosed by the Lon domain, the HB domain, the TBD and the previously disordered N-terminal belt, which wraps around the compact Lon-sensor loop-TBD unit, stabilising the closed conformation.^5^ New contacts also form between the TBD and HB domain. Cryo-EM studies suggest IMiD-bound CRBN exists in an equilibrium of the open and closed states.^5^ The bound IMiD creates a neosurface on CRBN, facilitating recruitment of neosubstrates such as GSPT1, CK1α, Ikaros, Aiolos, Helios and SALL4, for ubiquitination and subsequent degradation by the proteosome. IMiDs and other ligands that bind the tri-tryptophan pocket have been incorporated into PROTACs, expanding the neosubstrate target scope of CRBN. Neosubstrates can form protein-protein interactions with the Lon domain, TBD-sensor loop and N-terminal belt of CRBN. Lenalidomide (Revlimid) has been used as a treatment for multiple myeloma and other hematological cancers for several years, with next-generation molecules, such as iberdomide and mezigdomide, currently in clinical trials.^6–8^ These next generation IMiDs or CELMoDs (CRBN E3 ligase modulatory drugs), shift the open-closed equilibrium to greatly favour the closed state, which has been proposed as an explanation for the superior degradation exhibited in comparison to first-gen IMiDs.^5^ However, the molecular basis for this ligand-induced closure has not been fully elucidated. Moreover, it has remained unclear whether CRBN closure is a prerequisite for degradation and a key question for those seeking to exploit CRBN for targeted protein degradation.

Recruitment of CRBN has been extensively exploited for the development of PROTACs and molecular glue degraders, largely due to the availability of high affinity ligands, such as the IMiDs, and structural enablement facilitating structure-guided drug design.^3,4^ However, utilising IMiD binders to degrade proteins of interest safely via PROTACs while avoiding unwanted neosubstrate degradation poses a significant risk, usually requiring careful design and neosubstrate counter screening.^9^ Additionally, IMiDs have a racemisable stereocentre ^10,11^ and are prone to hydrolysis and ring opening at physiological pH,^12^ where the ring-open form can no longer bind to CRBN. Although there is a plethora of compounds approved and in clinical trials with this motif, various complex work packages are required during development due to the aforementioned challenges. Thus, several strategies have been employed to mitigate the associated liabilities, resulting in the development of alternative motifs such as dihydrouracils (DHUs) and glutarimides ^13–15^ Such strategies have been extensively documented for molecular glue degraders and PROTACs^16–21^ with several PROTACs now in clinical trials, for example, from C4 Therapeutics, Arvinas, BMS and Nurix Therapeutics.^22^ Despite this, there remains a need to identify novel CRBN binding motifs which may accelerate the speed with which promising degrader-based therapies reach patients.

To unlock novel chemical space and identify motifs capable of inducing targeted protein degradation, we initiated a project to identify novel CRBN binders. Using three different hit finding approaches, we discovered non-glutarimide scaffolds binding to CRBN as well as known CRBN-binding motifs. Two of these new series, morpholinone and spirolactam, were improved using rational design principles, guided by X-ray crystallography, to a suitable affinity range for PROTAC development. During the validation and characterisation of these molecules, we discovered that they behaved differently to known CRBN binders; IMiDs, dihydrouracils and glutarimides. Through extensive biophysical and structural characterisation of these new compounds, we found that they bound to the canonical tryoptophan cage pocket of CRBN but failed to induce CRBN closure. Thus, we were able to group the CRBN binders into two main classes: those that can induce CRBN closure, and those that cannot. Using mutational analysis, we also identified residues essential for compound-induced CRBN closure and show that mutation of these residues abrogates PROTAC-mediated degradation of BRD4 in cells by dBET6, a foundational IMiD-based PROTAC. Together, these data reveal insights into the CRBN open-closed equilibrium and imply that chemically induced CRBN closure strongly influences and may correlate with induced protein degradation activity by CRBN.

## Results

### Screening approaches to identify novel CRBN binders

To discover novel CRBN binders, we utilized a three-pronged approach consisting of a virtual screen (VS), high-throughput screening (HTS) and fragment-based screening (FBS) (Figure 2). In the VS workflow, we screened an internal compound collection with OpenEye’s shape-based tool ROCS (Rapid Overlay of Chemical Structures)^23^ using the thalidomide pose extracted from a published CRBN:thalidomide complex (PDB 7BQU). Hits were then docked back into the CRBN structure using Schrödinger’s Glide ^24^ Post-docking, poses were filtered using a pharmacophore (PH4) model which captured key interactions between thalidomide and CRBN (Figure 2A).^24^

**Figure 2.**
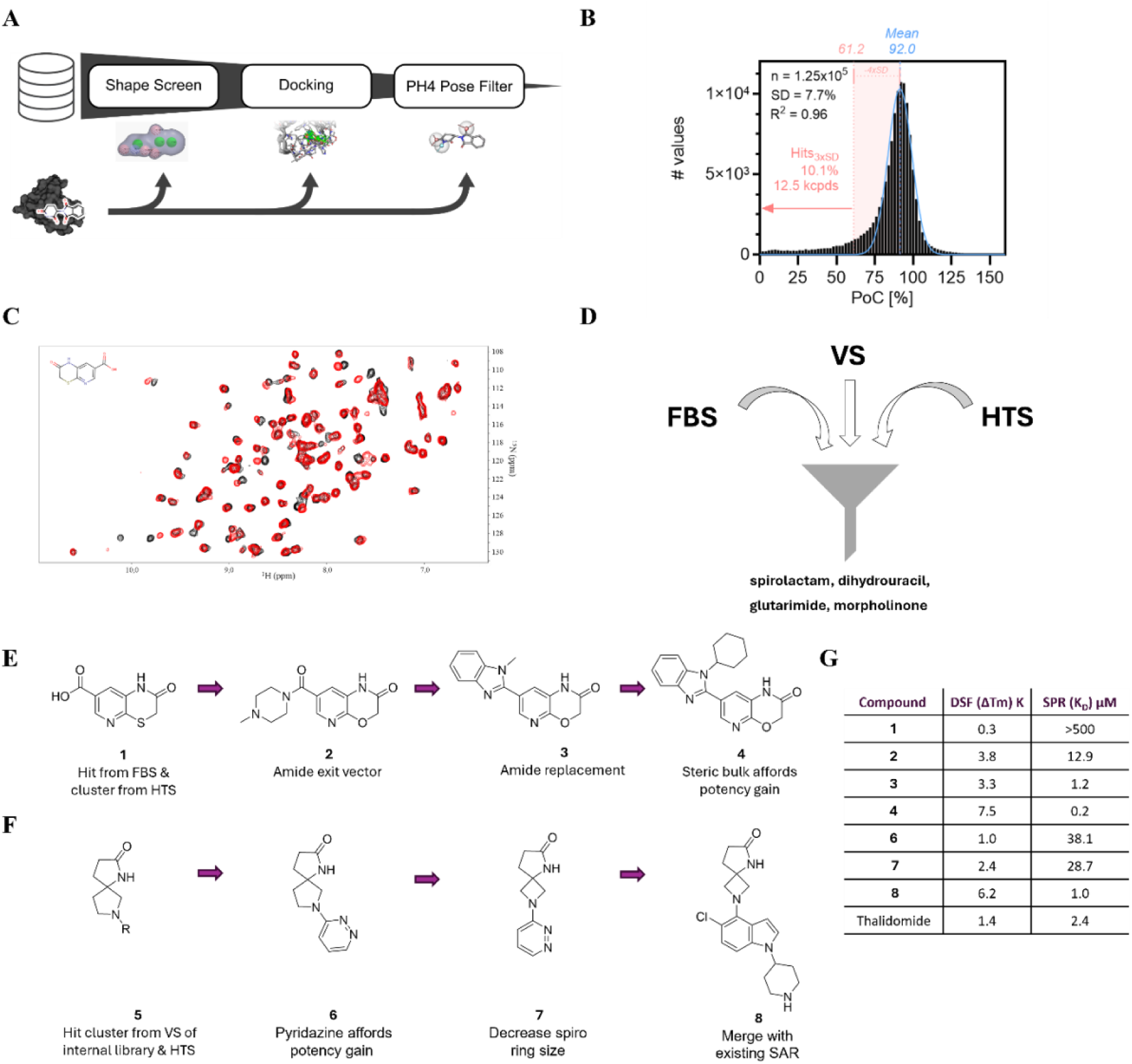
Identification and optimization of novel CRBN binders. Three-pronged screening strategy, deploying (A) virtual screening workflow (B) high-throughput screening of 12.5k compounds, using a TR-FRET-based reporter displacement assay with Cy-5 labeled Lenalidomide/His-tagged CRBN-DDB1 and (C) fragment-based screening utilizing protein observed NMR with 15N-labelled CRBN^TBD^ to identify novel CRBN binders, resulting in (D) the identification of four distinct chemical classes. E) Representative chemical structures of the morpholinone series optimisation resulting in high affinity compound **4**, F) Representative chemical structures of the spirolactam series optimization resulting in high affinity compound **8**, G) SPR affinity (K_D_) and DSF thermal shift data (ΔTm) for representative compounds using CRBN^TBD^.

We also performed an in-house high-throughput screen (HTS) as an unbiased approach to complement the biophysical fragment-based screening (FBS) and virtual screening (VS) methods. Using a proprietary library of low-MW molecules (<280 g/mol), compounds were screened at a single concentration in a TR-FRET-based reporter displacement assay designed to detect binding to the orthosteric CRBN tri-tryptophan pocket (Figure 2B).

In the FBS, we screened an in-house fragment library consisting of 8340 small molecules by protein observed NMR using a uniformly ^15^N labelled CRBN thalidomide binding domain (CRBN^TBD^) construct, since the molecular weight of full length CRBN*DDB1 exceeds the detectable size limit of NMR screening conditions. Mixtures experiencing significant chemical shift perturbation (CSP) were deconvoluted and yielded hits of mainly previously reported chemical matter such as glutarimides, dihydrouracils, uracils, hydantoins, and maleimides. In addition, less common core structures like morpholinones and thiomorpholinones were identified. The superposition of the 2D ^15^N HSQC of the latter with the DMSO reference spectrum is depicted in Figure 2C. The detected CSP is comparable to the CSP detected with thalidomide (Figure S1), indicating that the identified fragment hit binds to the same orthosteric interaction site formed by the tri-tryptophan cage.

Several hits identified by FBS and VS approaches were also found in the HTS, providing orthogonal validation and a strong foundation for further chemical optimization and characterization. Combined, these approaches afforded novel non-glutarimide CRBN binders, two of which, the morpholinone and spirolactam series, are detailed herein.

### Optimisation of Screening Hits

The first cluster identified by FBS & HTS were the morpholinones, as exemplified by FBS hit Compound 1 (Figure 2E). Utilizing homology modeling to the cocrystal structure of thalidomide and CRBN (PDB 7BQU), the 2- and 3-positions on the pyridine were identified as possible suitable exit vectors. The acid of compound **1** was therefore diversified by amidation and the sulfur of the lactam ring substituted with oxygen to afford compound **2**. This displayed increased affinity as measured by SPR (Figure 2G). Cyclization of the amide to a benzimidazole **3** improved affinity ten-fold. Increasing the steric bulk of the N-substituent on the benzimidazole to form compound **4** further increased affinity across all assays, ten-fold more potent than thalidomide.

The second cluster consisted of spirolactams with the common substructure exemplified by compound **5** (Figure 2F). As this motif alone did not display any measurable affinity to CRBN, it was substituted with different moieties, resulting in compound **6** with measurable affinity by SPR. Ring contraction to a 5,4 system afforded the spiro azetidine compound **7**, which displayed slightly improved affinity. Utilizing ligand-based modeling and structure overlays, the opportunity for elaboration based on known dihydrouracil CRBN binders was identified. As exemplified by compound **8**, this afforded a strong gain of affinity, bringing it into the same range as thalidomide. High affinity compounds **4** and **8** were taken forward for further characterisation.

### Use of CRBN constructs

To further characterise the new binder series and established binders, we employed two primary CRBN constructs. The isolated CRBN^TBD^ construct used in the NMR fragment screen and for initial DSF and SPR measurements is adequate to study direct ligand binding,^25^ but cannot recapitulate the allosteric rearrangement from the open to closed conformation that CRBN undergoes upon ligand binding.^5^ CRBN^TBD^ is also unsuitable for studying PROTAC-or molecular glue-induced ternary complexes, as these often involve neo-substrate binding surface interactions with the Lon domain of CRBN. Our recently reported CRBN^midi^ construct is soluble, expressed from *E. coli*, stable in the absence of DDB1 and contains the Lon, HB and TBD domains, with a short linker sequence replacing the DDB1 binding region between the Lon and HB domains.^26^ This construct is an important tool for biophysical and structural characterisation using techniques such as SPR, nano-DSF, SAXS and X-ray crystallography, with comparable functionality to the full-length wild-type CRBN. Importantly, CRBN^midi^ recapitulates the apo open and ligand-bound closed states of full-length CRBN. Therefore, once established, CRBN^midi^ became the construct of choice for biophysical experiments and X-ray crystallography over CRBN^TBD^ which had been used for the initial screening and optimization efforts. Full-length CRBN can be co-expressed with its cognate adaptor DDB1 in insect cells. DDB1 consists of three WD-40 beta propeller domains BPA, BPB, and BPC, with CRBN nestling into the hydrophobic cleft created at the junction between BPA and BPC. Whilst more challenging to make and work with, the CRBN:DDB1^ΔBPB^ (internal deletion of the flexible WD-40 beta-propeller B) complex is valuable for understanding the thermodynamics and induced allostery in relation to ligand binding. Therefore, both CRBN^midi^ and CRBN:DDB1^ΔBPB^ constructs were used for further structural and biophysical analysis of the CRBN binders.

### Comparison of different CRBN binder classes in biophysical assays

The binding affinity of the morpholinone and 5,4-spiro series was studied by SPR with the CRBN^midi^ construct, as the CRBN:DDB1^ΔBPB^ complex is larger and dissociates when immoblised under flow, making it unstable in SPR. C-terminally Avi-tagged CRBN^midi^ (C-avi-CRBN^midi^) was biotinylated and immobilised on a streptavidin chip and the binding of morpholinone **4** and 5,4-spirolactam **8** was measured and compared to example molecules from existing CRBN binder classes (Figure 3A and D, Table S1.). Compounds **4** and **8** had average steady state K_D_ values of 264 and 188 nM, respectively. K_D_ values in an equivalent range were obtained for lenalidomide (IMiD) and DHU **9** (K_D_ = 295 and K_D_ = 108 nM, respectively). This data supported that we had progressed the morpholinone and 5,4-spiro series into a similar affinity range as known binders that have been incorporated into PROTACs.^27^

**Figure 3.**
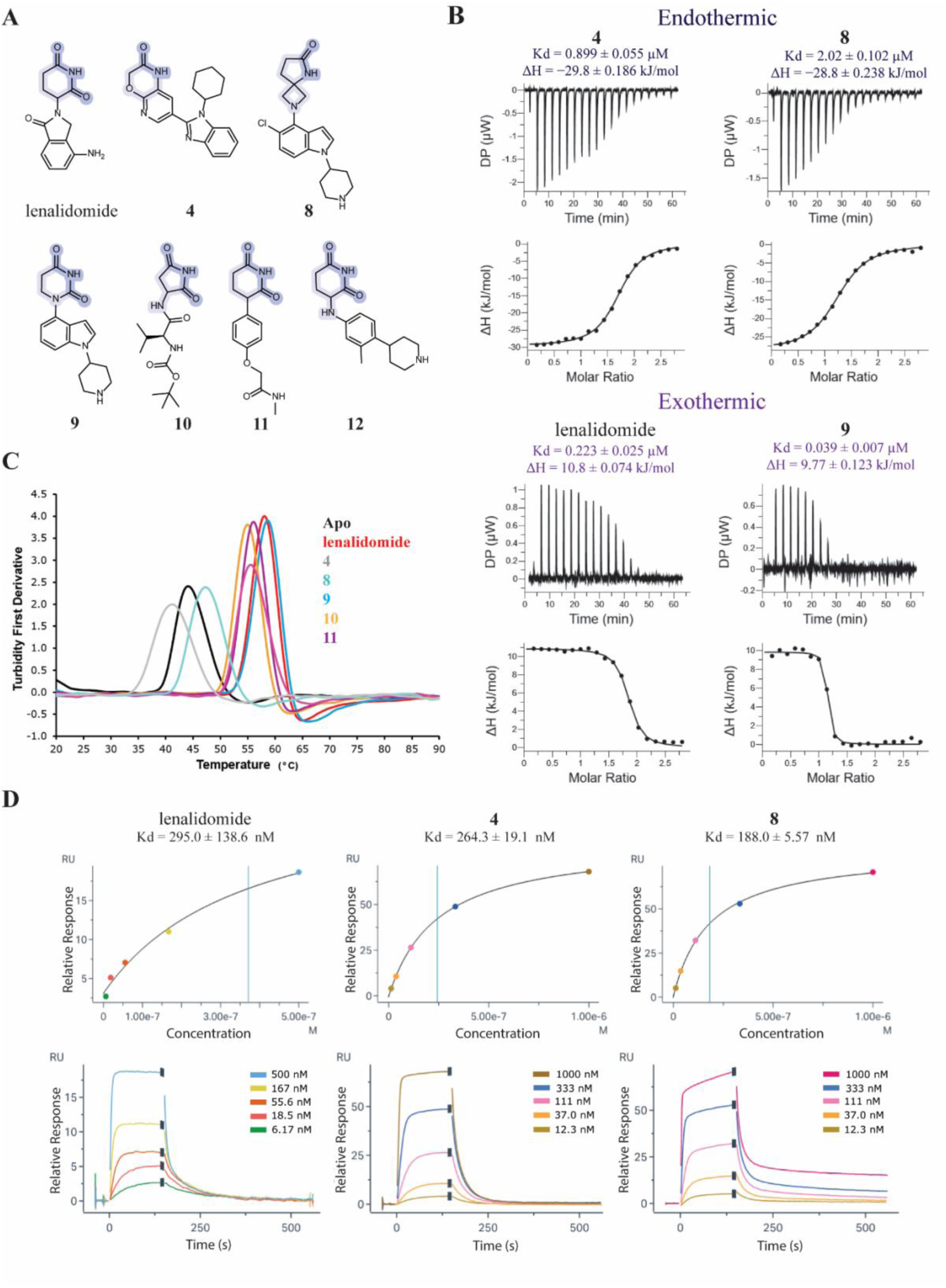
Biophysical binding studies in solution differentiate amongst binder classes. A) Molecular structures of CRBN binders measured across biophysics assays. Binding motifs are highlighted in blue; B) ITC measurements of selected CRBN binders with CRBN:DDB1^ΔBPB^, grouped into endothermic and exothermic binding; C) Turbidity first derivative of thermal denaturation for CRBN^midi^ in the absence (black) or presence of binders: lenalidomide (red), **4** (grey), **8** (aqua), **9** (blue), **10** (orange), **11** (purple) and **12** (pink). The average of three data points is shown; D) C-avi CRBN^midi^-on-chip SPR. Example sensorgrams and plotted steady state affinity graphs for lenalidomide, morpholinone **4** and spirolactam **8**.

We next used Isothermal Titration Calorimetry (ITC) as an orthogonal binding assay, utilising the full-length CRBN:DDB1^ΔBPB^ complex, and observed an interesting difference in the binding thermodynamics with the different binder classes. For lenalidomide (IMiD), dihydrouracil **9** and phenylglutarimide **11** binding to CRBN:DDB1^ΔBPB^ was observed to be endothermic, whereas binding of the newly discovered morpholinone **4** and spirolactam **8** was an exothermic process (Figure 3B, Table S2). These data pointed to a marked difference between our new binders and known CRBN chemical matter in their interaction with CRBN.

To further explore binding of different compound classes to CRBN, nano differential scanning fluorimetry (nanoDSF) was performed with CRBN^midi^. It has previously been demonstrated that lenalidomide significantly thermostabilizes CRBN^midi^ but not CRBN:DDB1.^26^ A potential explanation is that the melting temperature of the CRBN:DDB1 complex is dominated by the large and stable DDB1 protein, making CRBN^midi^ the more suitable construct for this technique. The melting temperature (T_m_) of apo CRBN^midi^ was 44.1°C and the thermal stability was substantially increased by between 10.9 to 14.6 °C with the addition of lenalidomide, DHU **9**, cyclimid **10**, phenylglutarimide **11** or aminoglutarimide **12** (Figure 3C, Table S3). Conversely, the addition of the 5,4-spiro **8** only increased the T_m_ by 3.2°C and the presence of the morpholinone **4** decreased the thermal stability of CRBN^midi^ with a ΔT_m_ of −2.9 °C. Measurements of additional compounds for the 5,4-spiro, morpholinone and DHU classes revealed series-specific thermostability shifts in the range of −2 to 4°C, −4 to 2°C, and 9 to 16°C, respectively (Table S3). The difference in the degree of thermostabilization by different compound classes is not explained by differences in binding affinity. For example, compounds **8** and **4** have relatively low SPR K_D_ values of ∼200 nM but have poor thermal stabilization of −2.9 and 3.2 °C. By contrast, compounds **10**, **11** and **12** have relatively high SPR K_D_ values (3.38 μM, 2.52 μM and 32.73 μM respectively) but have much higher ΔT_m_ values of >10 °C. In contrast to the results with CRBN^midi^ which contains both the Lon domain and TBD, ΔT_m_ values for CRBN^TBD^ alone were highest for the morpholinone and 5,4-spiro compounds (Table S3).

### Structural characterisation of ligand-induced CRBN closure

The striking differential behaviour of the new 5,4-spiro and morpholinone binder classes, exhibiting exothermic binding in ITC and lower thermostabilisation in DSF compared to established CRBN binders, led us to hypothesise that the new ligands were failing to induce CRBN closure. We therefore set out to structurally characterize binding of the different ligand classes using small-angle X-ray scattering (SAXS), X-ray crystallography and cryo-EM. SAXS coupled to in-line SEC can be used to analyse the rearrangement of the TBD and Lon domains of CRBN^midi^ induced by ligand binding.^26^ Apo CRBN^midi^ was found to have a radius of gyration (R_G_) of 26.83 Å, consistent with previous findings and the open conformation (Figure 4A).^26^ Addition of 400 μM ligand significantly reduced R_G_ with lenalidomide (22.99 Å), dihydrouracil **9** (23.43 Å) and phenylglutarimide **11** (22.8 Å), consistent with closed CRBN^midi^ and previous data obtained for the natural degron cyclimid scaffolds.^26^ Comparatively, the R_G_ for spirolactam **8** and morpholinone **4** aligned with the open conformation of CRBN^midi^, with values of 26.01 Å and 25.72 Å, respectively.

**Figure 4.**
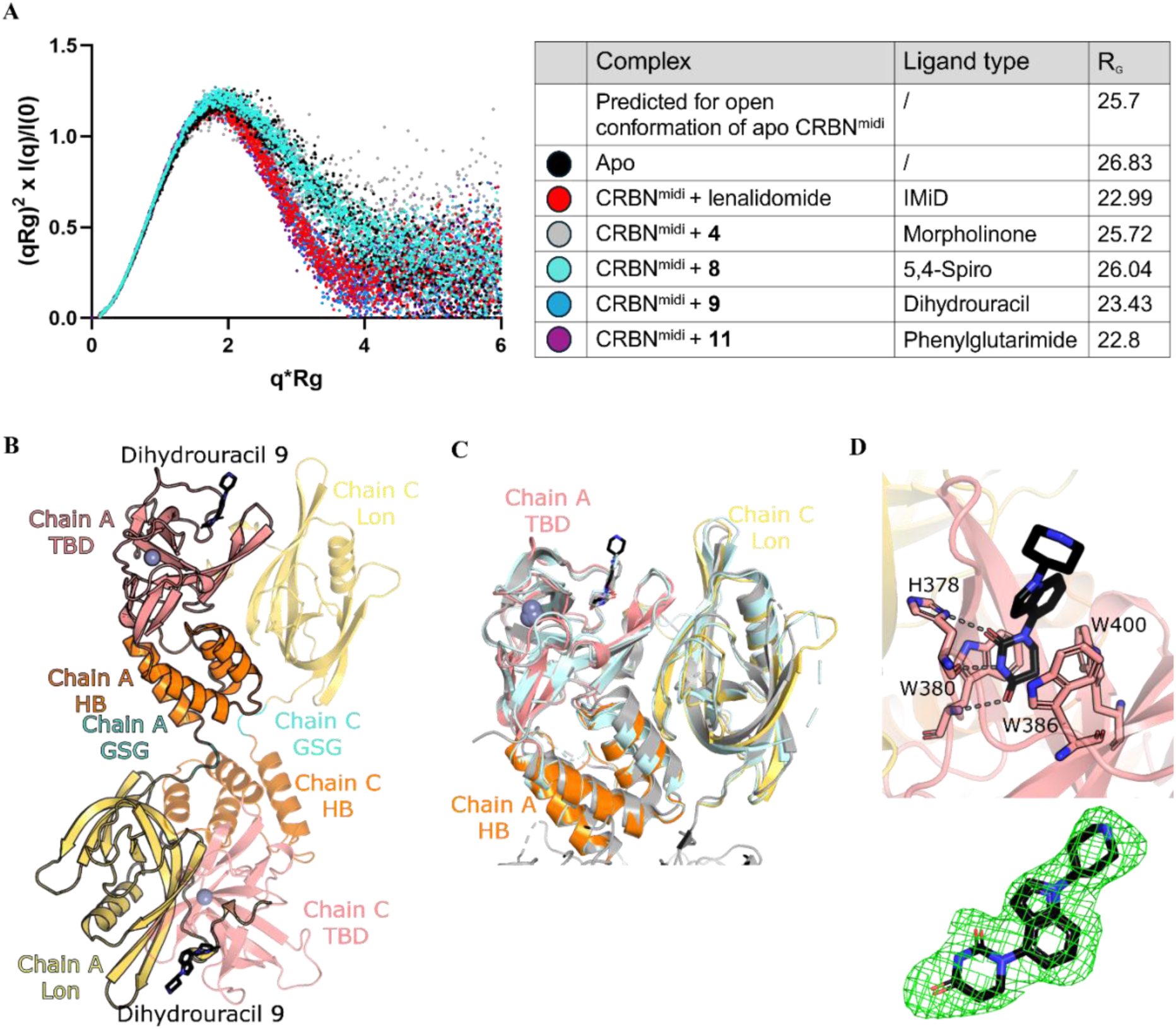
Structural comparison of different binder classes by SAXS and X-ray crystal structure of CRBN ^midi^ bound to dihydrouracil ligand. A) Dimensionless Kratky plot generated from SAXS data of Apo CRBN^midi^ (black) and CRBN^midi^ in complex with binders: lenalidomide (red), **4** (grey), **8** (cyan), **9** (blue) and **11** (purple). - Crystal structure of **9** bound to CRBN^midi^ and comparison to literature structures. B) Crystal structure of CRBN^midi^ in complex with compound **9**. CRBN^midi^ is shown in cartoon representation coloured by domain: Lon (yellow), HB (orange), TBD (salmon), GSG linker (cyan). **9** is shown as black sticks and Zn^2+^ as a grey sphere. **a** Chain A is outlined in black to show the domain swap arrangement. C) superposition of Lon (chain C), HB and TBD (chain A) domains of CRBN^midi^:**9** with the crystal structure of CRBN^ΔN^:DDB1:lenalidomide (grey, PDB ID: 4TZ4) and CRBN^midi^:lenalidomide (pale cyan, PDB ID: 8RQA). D) TBD binding site with **9** (chain A). Trp cage residues and H378 are shown in stick form, hydrogen bonds are represented by dashed grey lines. Polder OMIT (Fo-Fc) map of the ligand contoured to 3 σ in each chain shown as green mesh.

We next sought to visualise the binding mode of the new series and dihydrouracil **9** using X-ray crystallography. Numerous binary crystal structures of CRBN in complex with thalidomide-based scaffolds are present in the PDB with CRBN:DDB1, CRBN^midi^ and CRBN^TBD^ (PDB IDs: 9FJX, 8OIZ, 8OJH, 4CI3, 4CI1, 4CI2, 4TZ4, 5V3O, 8G66, 8RQA, 8RQ8, 9ODR, 9ODR). The recently solved structure of CRBN^midi^ in complex with phenylglutarimide **11** (PDB ID: 9GAO) revealed an identical binding mode of the glutarimide ring to lenalidomide.^26^ Excluding CRBN^TBD^ which lacks the Lon and HB domains, CRBN adopts the closed conformation in all these binary crystal structures. CRBN^midi^ was co-crystallised with dihydrouracil **9**^27^ and the structure was solved to 2.36 Å (Figure 4B), to our knowledge the first reported structure of CRBN with a dihydrouracil ligand. During model building, a domain swap was observed between the Lon domain at one side of the GSG linker and the HB/TBD domains on the other (Figure 4B). Electron density was observed for the GSG linker in both the domain swapped and non-domain swapped organisations (Figure S2A), suggesting that this phenomenon may occur heterogeneously throughout the crystal. The refinement statistics were improved with the domain swap; hence it was chosen as the final model. The biologically relevant Lon-HB-TBD domain unit of the CRBN^midi^:**9** structure adopts the same overall fold as previous CRBN:lenalidomide structures in the closed conformation (Figure 4C). Superposition of CRBN^midi^:lenalidomide (PDB ID: 8RQA) and CRBN^Δ1–40^:DDB1:lenalidomide (PDB ID: 4TZ4) yielded RMSDs of 0.633 Å and 1.088 Å, respectively. The binding mode of dihydrouracil **9** is comparable to the glutarimides and an uracil-containing CRBN binder (PDB ID: 9ODS, Figure S2B),^28^ maintaining the tripartite hydrogen bonding interactions with the backbone carbonyl and side chain of H378 and the backbone nitrogen of W380 (Figure 4D). Co-crystallisation of CRBN^midi^ with morpholinone and spirolactam compounds **4** and **8** was attempted, however, crystals that diffracted to a resolution high enough to determine the structures could not be generated, consistent with the hypothesis that these binders fail to stabilise the closed conformation of CRBN.

To further investigate the influence of compound binding on the structural dynamics of CRBN, we determined the cryo-EM structures of the dihydrouracil ligand **9** and the spirolactam ligand **8** bound to CRBN:DDB1^ΔBPB^ (Figure 5, Figures S3-7, Tables S8-9). Previously solved cryo-EM structures of CRBN bound to pomalidomide exhibited an equilibrium of closed and open CRBN.^5^ Consistent with this, we observed CRBN in both the closed (Figure 5A) and open conformation (Figure 5B) with DHU **9**, refining reconstructions of both conformations to a global resolution of 3.33 and 3.66 Å, respectively (see Figure S4 for consensus maps). Local refinement of TBD in the open conformation and Lon-TBD unit in the closed conformation yielded maps of 4.23 and 4.07 Å, respectively, with improved map quality around these regions of interest (see Figure S4 for locally refined maps). In the open conformation, we observed signal for the sensor loop adjacent to the compound (Figure 5B), suggesting engagement and ordering of the sensor loop follows ligand binding but precedes CRBN closure.

**Figure 5.**
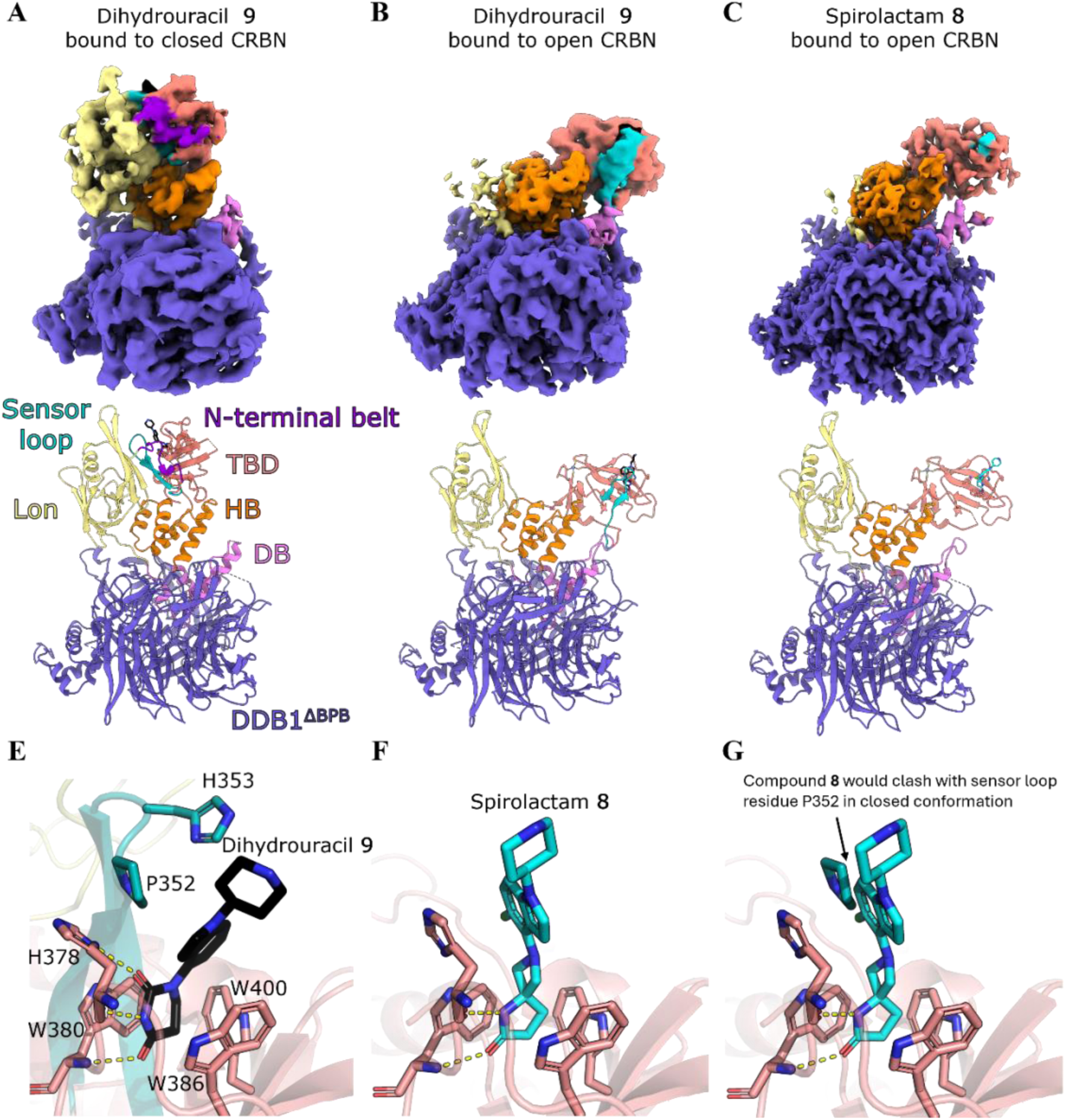
Structural comparisons of ligand binding to open vs closed CRBN. Composite cryo-EM maps and models of CRBN:DDB1^ΔBPB^ bound to dihydrouracil **9** in the closed (a) and open conformation (b), and 5,4-spiro **8** in the open conformation (c). The N-terminal belt, Lon domain, DDB1 binding (DB) region, Helical bundle (HB), thalidomide binding domain (TBD), sensor loop of CRBN and DDB1^ΔBPB^ are labelled in (a). (d) Binding of dihydrouracil **9** to closed CRBN in the CRBN^midi^ crystal structure. (e) Binding of spirolactam **8** to CRBN TBD.(f) Sensor loop residue P352 from closed conformation would clash with the chloroindole of compound **8**.

In the 2.22 Å reconstruction of CRBN:DDB1^ΔBPB^ with spirolactam **8** bound (Figure 5C), we observed only the open conformation and a disordered sensor loop, despite map signal for the ligand in the 3.44 Å locally refined map (Figure S6) of the CRBN TBD tri-tryptophan pocket (Figure S7). In comparison to dihydrouracil **9** (Figure 5D), spirolactam **8** makes two of the three canonical hydrogen bonds in the tryptophan pocket: to the backbone carbonyl of H378 and the backbone nitrogen of W380 (Figure 5E). The positioning of the chloroindole moiety differs significantly to the indole of compound **9**, potentially forming a π-stacking interaction with the sidechain imidazole ring of H378 (Figure 5E). In closed structures of CRBN bound to IMiDs, phenylglutarimide **11** and dihydrouracil **9**, the H378 imidazole instead acts as a hydrogen bond donor to the ligand, and forms van der Waals stacking interactions with sensor loop residue P352 (Figure 5D). In the closed conformation, P352 would clash with the chloroindole of the spirolactam **8** (Figure 5F), which may explain why spirolactams fail to induce CRBN closure, through steric blocking of sensor loop engagement. The morpholinone compound class lacks the second hydrogen bond acceptor found in other established CRBN binder classes (Figure 3A), which could also affect the conformation of H378 and therefore CRBN closure. We collected cryo-EM data for CRBN:DDB1^ΔBPB^ with a morpholinone compound and observed only the open conformation on 2D and 3D classes (data not shown). Taken together, our biophysical and structural data suggest that compound binding alone is not sufficient to induce CRBN closure, and that specific chemical features such as the tripartite hydrogen bonding motif and compound geometry compatible with the closed conformation may be required.

### Investigating CRBN closure

Given the apparent importance of CRBN closure for protein degradation, we set to identify key residues whose mutations could impair CRBN closure. To this end, we explored interactions around the ligand binding site and between the N-terminal belt, Lon domain, TBD and sensor loop, as observed in the closed state. Analysis of the 5FQD structure (CRBN:DDB1^ΔBPB^:lenalidomide:CK1α) revealed a network of hydrogen bonds and hydrophobic interactions involving Y59 and L60 (N-terminal belt), Q100 and K156 (Lon domain), V350 and N351 (sensor loop), and H378 and W380 (TBD) (Figure 6A and B).

**Figure 6.**
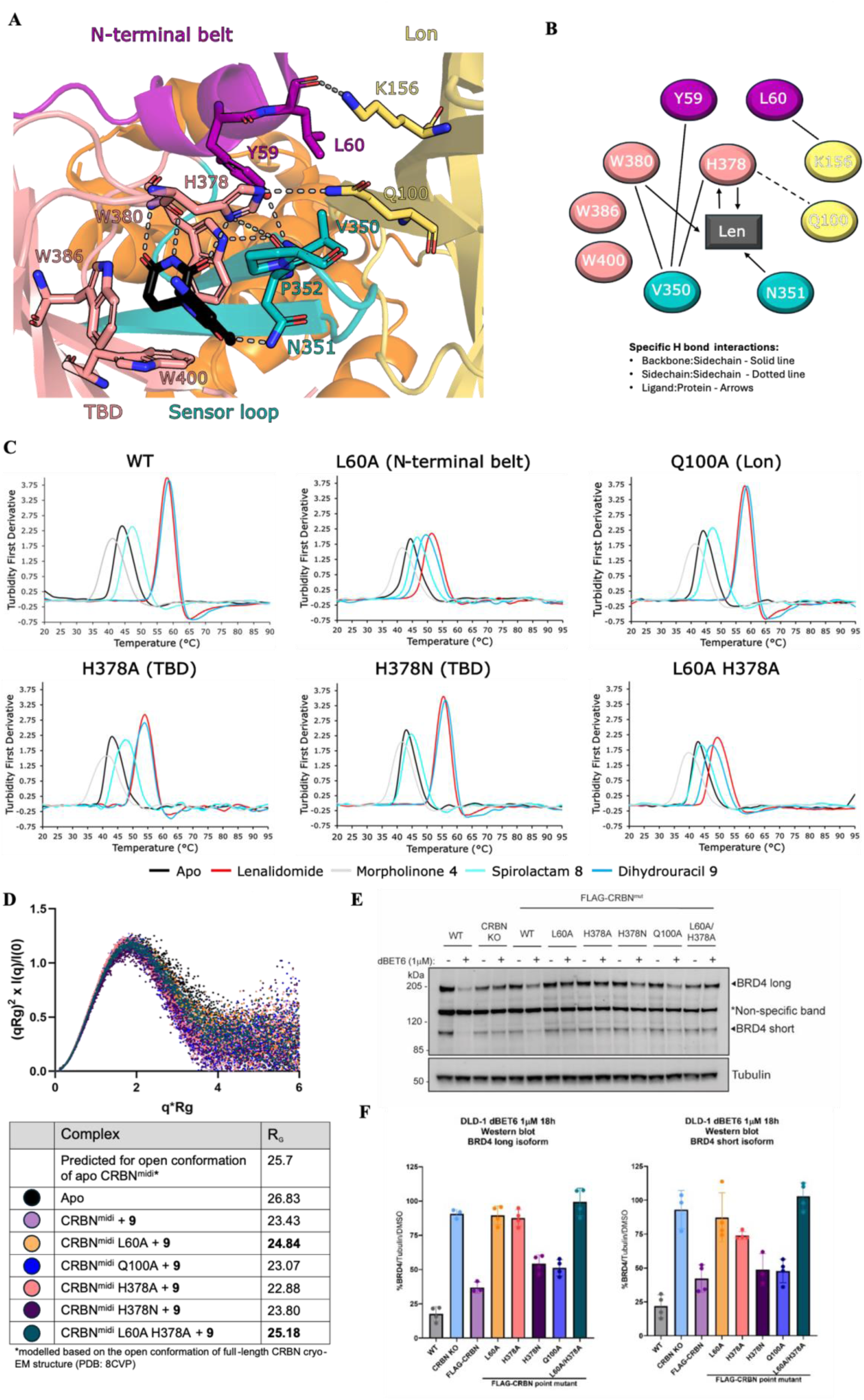
Residues L60 and H378 are critical for ligand-induced CRBN closure and PROTAC-dependent degradation. A) Structural analysis of closed CRBN and lenalidomide, showing the network of hydrogen bonds in 5FQD. The structure is coloured by domain Lon (yellow) (N-terminal belt (purple), HB (orange), TBD (salmon), (sensor loop (teal)). Lenalidomide is shown as black sticks, the Tryptophan cage and residues involved in hydrogen bonds are shown in stick form, hydrogen bonds are represented by dashed grey lines. B) A schematic representation of the network of interactions. Backbone to sidechain H-bonds are represented by solid lines, sidechain to sidechain H-bonds by dotted lines and ligand to protein H-bonds by arrows. C) Turbidity first derivative of thermal denaturation for CRBN^midi^ WT and mutants in the absence (black) or presence of binders: lenalidomide (red), **4** (grey), **8** (cyan) and **9** (blue). WT is mean of triplicates. D) Dimensionless Kratky plot and table of Rg values generated from SAXS data of CRBN^midi^ and CRBN^midi^ mutants in complex with dihydrouracil **9**. E) Representative Western Blot assessing BRD4 degradation efficiency in DLD-1 WT cells, DLD-1 CRBN KO cells and DLD-1 CRBN KO cells stably expressing FLAG-CRBN WT or CRBN point mutants. Cells were treated with dBET6 (1 µM) for 18h and degradation of BRD4 long and short isoform was quantified and normalised against a loading control and DMSO-treated samples. F) Quantification of the bands from (E).

To investigate the importance of these residues for CRBN closure, we attempted to generate single point mutants of CRBN^midi^, covering at least one residue from each section of the network. The N-terminal belt residues L60 and Y59 bury their side chains in a hydrophobic pocket formed by the Lon domain, sensor loop and TBD, so we attempted to mutate both residues to alanine. The side chain of Q100 in the Lon domain forms a hydrogen bond with the ligand binding site residue H378 in the TBD, so we mutated this to alanine. In addition to the interaction with Q100, the H378 side chain is involved in hydrogen bonding interactions with the glutarimide ring of lenalidomide and the backbone carbonyl of V350 in the sensor loop. We mutated H378 to alanine or asparagine, which would remove the hydrogen bonding interactions or impair stacking with P352 in the sensor loop, respectively. We also attempted to mutate V350 in the sensor loop to glutamate to disrupt interactions with the Lon domain. Single point mutants were successfully generated in CRBN^midi^ for L60A, Q100A, H378A and H378N, which expressed and purified comparably to wild-type, while Y59A and V350E failed to express any soluble protein.

The impact of the point mutations on CRBN^midi^ thermostabilisation was investigated by nanoDSF in the presence of lenalidomide, dihydrouracil **9**, spirolactam **8**, and morpholinone **4** (Figure 6C, Table S4). Thermostabilisation of CRBN^midi,Q100A^ and CRBN^midi,H378N^ was comparable to wild type all binder classes, suggesting that the loss of a hydrogen bond between residues Q100 and H378 and the loss of stacking potential with P352 does not impact the ability of CRBN to close. CRBN^midi,H378A^ showed reduced thermostabilisation by lenalidomide and dihydrouracil **9** compared to CRBN^midi^, with a reduction in ΔT_m_ of 3 - 4 °C. In contrast, the thermostabilisation of CRBN^midi,H378A^ by morpholinone **4** and spirolactam **8** increased ∼1.5°C compared to CRBN^midi^. This could be explained by the loss of the hydrogen bond from the side chain of H378 to lenalidomide and the dihydrouracil **9** that is not present with compounds **4** and **8**. However, the H378A mutation has previously been shown to not impact the binding of IMiDs to CRBN^TBD^.^29^ The most significant change was observed with the L60A mutation in the N-terminal belt. Compared to CRBN^midi^, a large reduction in ΔT_m_ was observed with CRBN^midi,L60A^ for closure-inducing compounds lenalidomide and dihydrouracil **9** (ΔT_m_ = 6.9°C and ΔT_m_ = 9.35 °C, respectively), while the ΔT_m_ for morpholinone **4** and spirolactam **8** remained consistent with that observed for CRBN^midi^. The double mutant CRBN^midi,L60A/H378A^ was generated to test if there was an additive effect for the reduction in thermostabilisation. The double mutant showed similar ΔT_m_ values to the single CRBN^midi,L60A^ mutant, suggesting the additional loss of H-bonding from the H378 mutation did not further exacerbate the defect seen with L60A.

To confirm the reduction in the degree of thermostabilisation of CRBN^midi,L60A^ and CRBN^midi,L60A/H378A^ by dihydrouracil **9** was due to impaired CRBN closure, we carried out SAXS with each of the mutants in the presence of 400 μM dihydrouracil **9** (Figure 6D). CRBN^midi,L60A^ and CRBN^midi,L60A/H378A^ were both found to have increased R_G_, consistent with impaired capacity of adopt the closed conformation. We refer to these mutants as “closure-deficient”. The remaining mutants all exhibited radii of gyration similar to CRBN^midi^, consistent with the closed conformation and results from DSF experiments.

### Investigating the impact of closure-deficient CRBN on degradation

To assess the effect of CRBN mutations on PROTAC degradation efficiency, six stably expressing cell lines were generated in a DLD-1 CRBN knock-out (KO) cell line, expressing FLAG-tagged: CRBN^WT^, CRBN^L60A^, CRBN^H378A^, CRBN^H378N^, CRBN^Q100A^ and CRBN^L60A/H378A^. Each cell line was then assessed for PROTAC-induced degradation of BRD4 by treatment with the potent CRBN-based BET degrader dBET6 (Figure 6E and F).^30^ In the parental DLD-1 CRBN KO cell line, degradation of both long and short isoforms of BRD4 by dBET6 was abrogated, as expected. Re-expression of FLAG-tagged CRBN^WT^ led to incomplete rescue of BRD4 degradation, with a D_max_ of approximately 60% following 18 h incubation with 1 µM dBET6. In comparison, treatment of WT DLD-1 cells with dBET6 led to a D_max_ of approximately 80% at the same timepoint. We hypothesised this could be due to some instability in the re-expressed FLAG-tagged version of CRBN. To test the stability of FLAG-CRBN, we performed a time course experiment with WT DLD-1 cells endogenously expressing WT CRBN and DLD-1 CRBN KO cells stably expressing FLAG-tagged CRBN^WT^, treating with cycloheximide to inhibit protein synthesis (Figure S8). Unlike wild-type CRBN, the FLAG-tagged CRBN levels decrease over the eight-hour experiment, indicating a loss of protein stability which explains the lower-than-expected D_max_ levels.^31^

In agreement with our biophysical and SAXS data, neither long nor short isoforms of BRD4 were degraded in cells stably expressing CRBN closure-deficient mutants CRBN^L60A^ and CRBN^L60A/H378A^ when treated with dBET6 (Figure 6E and F). dBET6 also failed to degrade BRD4 in the cell line expressing CRBN^H378A^, which showed moderately reduced thermostabilisation by lenalidomide and dihydrouracil **9** but radius of gyration consistent with closed CRBN^midi^. This could be due to a reduction in affinity related to the loss of a H-bond in the tripartite H-bonding motif. Cells expressing CRBN^WT^, FLAG-tagged CRBN^WT^ and “closure-competent” mutants CRBN^H378N^ and CRBN^Q100A^ competently degraded of both BRD4 isoforms upon treatment with dBET6 (Figure 6E and F).

The L60 residue in the N-terminal belt does not directly interact with binders and is far distal to the binding site in the open conformation of CRBN, disconnecting its effect from small molecule binding to the orthosteric tri-tryptophan pocket. Our data suggests the loss of the hydrophobic leucine side chain disrupts the allosteric effect of the N-terminal belt wrapping around the Lon-TBD unit in the closed conformation, thereby impacting the ability of CRBN to close. This finding is consistent with previous work showing that deletion of the N-terminal belt abolishes CRBN closure observed by cryo-EM^5^ and demonstrates for the first time that CRBN closure is required for ubiquitination and degradation.

## Discussion

Small-molecule CRBN ligands typically contain a glutarimide functionality that closely mimics the native cyclic imide C-terminal degron recognized by CRBN.^32^ These ligands, such as IMID molecular glue degraders lenalidomide and mezigdomide, have been found to stabilize CRBN into a closed state.^5^ However, the extent to which the open-close equilibrium influences neo-substrate degradation by CRBN in cells has remained unclear. We compared and contrasted the behaviours observed for different CRBN binder classes in the biophysical and structural techniques used in this study (Table 1). We observe a striking consistency with essentially all methods converging to categorize between ligand classes that induce CRBN closure and those which do not. We show that DSF, SAXS and ITC alone can distinguish between the two binders categories. These solution-based biophysical methods are less resource and time intensive than cryo-EM or crystallography and can be used to prioritise ligands that stabilize the closed conformation. The binding assays presented herein will therefore enable quick assessment of whether new CRBN binders can induce CRBN closure and to what extent, and therefore inform on their likelihood to generate active PROTACs or molecular glue degraders.

**Table 1.**
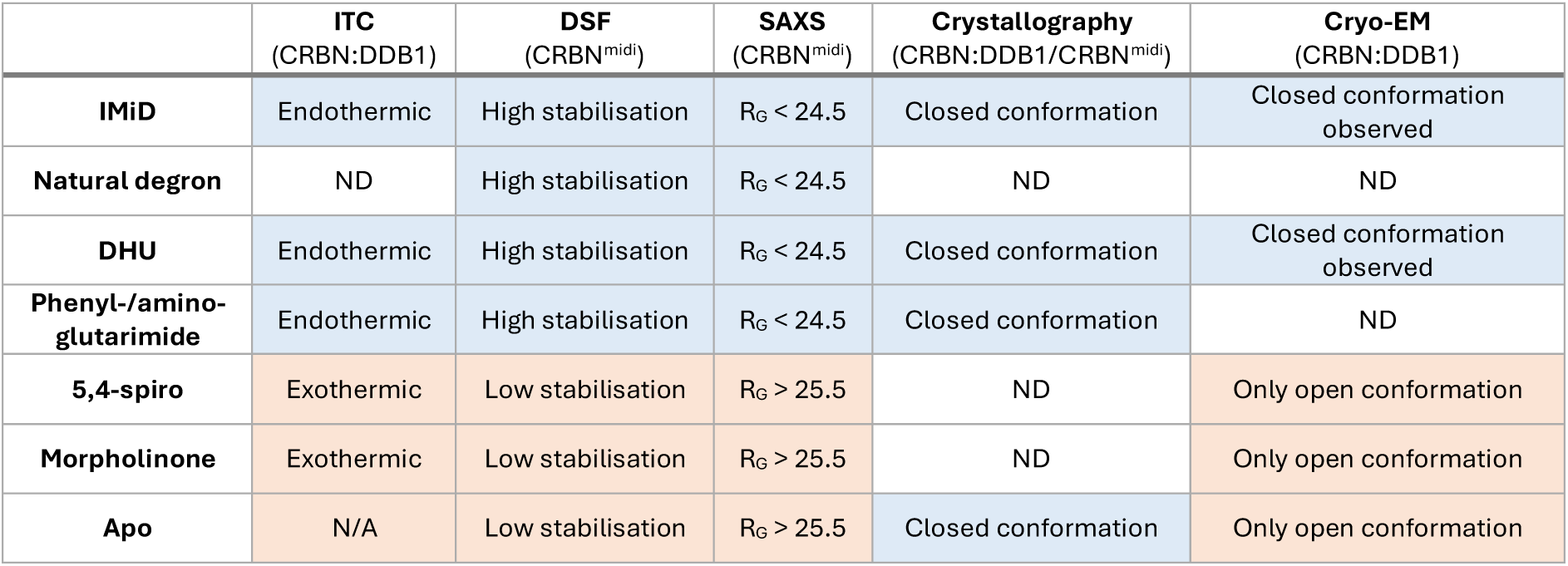
Summary of assay results for different CRBN binder classes.

The well-characterized native degron and IMID ligands, as well as the second-generation DHUs and glutarimide ligands, now widely used in both molecular glues and PROTACs, are all able to close CRBN. Contrasting these, we show that our pharmacophoric binder chemotypes of 5,4-spiro and morpholinones still bind to the orthosteric tri-tryptophan pocket in the TBD, but are unable to close CRBN. This class of binders will be expected to bias the open state of CRBN in the cell. They could therefore be utilized as chemical tools to explore biological and structural aspects of open vs. closed CRBN, and to probe the functional consequences of inhibiting CRBN without any confounding effects from CRBN-mediated degradation of neo-substrates. To accelerate efforts in these directions, we have identified and qualified a high-affinity morpholinone ligand **4** that, together with its negative control N-methylated analogue, will be made available on the Boehringer Ingelheim OpnMe platform free of charge.^33^

Understanding how different classes of CRBN binders affect the dynamics of CRBN conversion from open to closed states is important to guide more rational design of chemical probes and drugs targeting CRBN. Through structural characterization by cryo-EM and reconstitution of closure-deficient CRBN mutants in cells, we identify an intricate network of inter- and intra-domain interactions in addition to protein-ligand interactions that together orchestrate the open-close equilibrium. We find that ligands that are able to close CRBN are distinct from those that do not due to their ability to form a critical hydrogen bond with the side chain of TBD domain residue H378, reorienting it to “recruit” key residues within the otherwise disordered sensor loop and to recruit the Lon domain. These interactions create an interfacial pocket where the N-terminal belt latches into, filling the pocket with its L60 side chain. Mutations of H378 and L60 were found to impair CRBN closure, and to abrogate CRBN-dependent Brd4 protein degradation by dBET6 as archetypical representative of CRBN-recruiting PROTAC. Together with our inability to demonstrate induced protein degradation with any of the PROTACs we have made so far using our novel, closure-incompetent CRBN binder chemotypes, we conclude that effective targeted protein degradation via CRBN is greatly enhanced (and potentially only possible) when the E3 ligase receptor is in its closed state. Looking forward, the molecular and structural insights revealed in this study on the open-closed equilibrium and how this process can be fine-tuned chemically and with missense mutations, are anticipated to impact the design of state-specific tools to study CRBN biology and to help innovate next-generation of CRBN-targeting agents.

## Acknowledgements

We acknowledge the technical and research staff of the Ciulli Laboratories at CeTPD for the set-up and upkeep of protein expression and purification infrastructure. We would like to thank Maria Rodriguez-Rios for providing us with BRD4 BD2 protein for SPR experiments, and Roberta Ibba for helping with intact protein mass spec analysis (Ciulli Group, CeTPD, University of Dundee), and the research and lab management staff at CeTPD for technical support. We also thank Prof. Gopal Sapkota (MRC PPU, University of Dundee) for providing us with the DLD-1 WT and CRBN KO cell lines. We thank the Diamond Light Source for beamtime (BAG proposals SM38813 and SM42296 for SAXS) and the B21 team and Katsuaki Inoue at Diamond for running the SAXS samples. The University of Dundee Cryo-EM facility is funded by the Wellcome Trust (223816/Z/21/Z) and the MRC (MRC World Class Laboratories PO 4050845509). Molecular graphics and analyses were performed with both PyMOL (The PyMOL Molecular Graphics System, Version 2.5 or 3.0, Schrödinger, LLC) and UCSF ChimeraX (developed by the Resource for Biocomputing, Visualization, and Informatics at the University of California, San Francisco, with support from National Institutes of Health R01-GM129325 and the Office of Cyber Infrastructure and Computational Biology, National Institute of Allergy and Infectious Diseases).

## Author Contributions

T.T. & S.O’C wrote the paper. Z.R. wrote the paper and conducted molecular biology, protein purification, biophysical and structural biology experiments. A.D.C. wrote the paper and solved Cryo-EM structures. S.O’C, M.K., F.B. & T.T. designed compounds & synthetic routes. L.M., G.K., G.P., M.S., and S.O’C. designed synthetic routes and synthesized compounds. E.M, I.P., L.C., E.H., Y.C. and L.S. created cell lines and designed and performed *in cellulo* degradation experiments. S.C. conducted computational analysis of CRBN. S.D. conducted DSF experiments. Q.Z. contributed ideas towards synthetic routes. V.V. performed early biological experiments aiding compound selection. M.Z., S.L., F.B. & T.T. conceived, supervised & evaluated the CRBN screening. A.Ci. and K.M. wrote the paper, designed experiments, and supervised the project.

## Competing interests

M.Z., L.G., S.L. and T.T. are current employees of Boehringer Ingelheim. M.K. is a current employee of Merck KGaA. A.Ci. is a scientific founder and shareholder of Amphista Therapeutics, a company that is developing targeted protein degradation therapeutic platforms, and is on the Scientific Advisory Board of ProtOS Bio and TRIMTECH Therapeutics. The Ciulli laboratory receives or has received sponsored research support from Almirall, Amgen, Amphista Therapeutics, Boehringer Ingelheim, Eisai, Merck KGaA, Nurix Therapeutics, Ono Pharmaceuticals and Tocris-Biotechne.

## Data Availability

All atomic coordinates and structure factors for X-ray crystallography have been deposited in the Protein Data Bank with accession code 9SUN. Cryo-EM coordinates have been deposited in the Protein Data Bank with accession codes 9SVG (Spiro), 9SVH (DHU, closed conformation) and 9SVI (DHU, open conformation) and maps to the Electron Microscopy Data Bank with accession codes EMD-55257 (Spiro, composite map), EMD-54624 (Spiro, consensus map), EMD-54644 (Spiro, local TBD map), EMD-55258 (DHU, closed composite map), EMD-55057 (DHU, closed consensus map), EMD-55058 (DHU, closed local LON-HB-TBD map), EMD-55259 (DHU, open composite map), EMD-55059 (DHU, open consensus map), and EMD-55060 (DHU, open local TBD map). SAXS data have been deposited to Small Angle Scattering Biological Data Bank (SASBDB) under accession numbers: XXXX. Experimental procedures and supplementary tables and figures can be found in the Supporting Information.

## Declaration

For the purpose of open access, the corresponding author has applied a Creative Commons Attribution (CC BYSupportinguthor Accepted Manuscript version arising from this submission

## Methods

### Molecular biology

CRBN^midi,L60A^, CRBN^midi,Q100A^, CRBN^midi,H378A^, CRBN^midi,H378N^ and CRBN^midi,L60A/H378A^ were generated by site-directed mutagenesis. For the single point mutations WT CRBN^midi^ was used as a template and for CRBN^midi,L60A/H378A^ the H378A mutation was introduced using CRBN^midi,L60A^ as a template. In each reaction, 50 ng of plasmid template was used with 100 ng of both the forward and reverse primers. 25 μL of Q5® High-Fidelity 2X Master Mix (New England Biolabs) was added and the reaction was made up to 50 uL using nuclease free water. All reactions were carried out using a ProFlex PCR System (Thermo Fisher Scientific). Mutagenesis products were analysed on a 1% agarose gel and parental DNA was digested with 16 U of *Dpn*I (New England Biolabs) for 2 hours at 37 °C followed by denaturation of *Dpn*I at 80 °C for 20 minutes. Successful mutagenesis products were transformed into DH5α cells and plated onto LB KAN agar plates. 5 mL of LB KAN was inoculated with a single colony and grown overnight at 37 °C. The plasmid was extracted and purified from the cells using GeneJET Plasmid Miniprep Kit System (Thermo Fisher Scientific). All DNA sequencing was carried out by MRC PPU DNA Sequencing and Services (University of Dundee) using universal T7 primers (T7 and T7 Term). A list of primers and cycling parameters can be found in Tables S10-11. All primers were manufactured by Merck (Sigma-Aldrich).

### Protein expression and purification

#### CRBN^midi^ and CRBN^TBD^

##### Stable isotope labelled CRBN^TBD^ for NMR studies

The DNA sequence of human Cereblon (Uniprot acc. no. Q96SW2, AA 318-426 with GST-Tag and TEV cleavage site between the tag and the protein of interest) was cloned into pETDest vector (Invitrogen) and transformed into BL21(DE3) cells (Invitrogen). Cells were grown in 15N OD2 medium (Silantes) supplemented with Ampicillin (30 μg/mL) until the OD (600 nm) reached 0.6. Protein expression was induced with 1 mM IPTG, the cells were cultivated for additional 16 h at 20°C, harvested by centrifugation and stored at -70°C.

For protein purification, the bacterial cell pellet (8 L culture) was thawed, suspended in lysis buffer (50 mM Tris-HCl, pH 7.5, 500 mM NaCl, 1 mM TCEP, Complete Protease Inhibitor Cocktail (Roche) and 1 mg DNAse) and disrupted by sonification. The crude cell lysate was clarified by centrifugation at 45000g for 60 min. The supernatant containing GST-tagged Cereblon was purified on 15 ml GT-Sepharose-Slurry (GE Healthcare). The protein was eluted by the addition of 4.5 mg TEV Protease (incubation over night at 4°C). The elution was further purified on a Superdex 75 column (Cytiva) in 20 mM Tris pH 7.5, 200 mM NaCl and 1 mM TCEP and concentrated to 2,5 mg/mL using a centrifugal filtering device (Millipore, 3 kDa molecular-weight cutoff).

CRBN^midi^ and CRBN^TBD^ proteins were expressed and purified as described previously^26^ with the addition of 10 % glycerol (v/v) in the affinity chromatography buffers. Point mutations to CRBN^midi^ were confirmed by intact protein LC-MS. A 10-75 % gradient of acetonitrile was used to separate the samples by HPLC (C3 column) they were then analysed using an Agilent 6130 quadrupole MS. Specta were deconvoluted and integrated using Agilent LC/MSD ChemStation, predicted and experimental masses are in Table S12.

##### Biotinylated C-avi-CRBN^midi^

N-terminally His_6_-tagged and C-terminally Avi-tagged CRBN^midi^ (C-avi-CRBN^midi^) was expressed in *E. coli* BL21 (DE3) cells. Cell cultures were grown in LB liquid medium at 37°C with 50 μg/mL kanamycin to an OD_600_ 0.7-0.9. Expression of the target protein was induced with 0.5 mM IPTG, 50 μM ZnCl_2_ was added to the media and the cultures were incubated at 18 °C for 20 hours. The cells from 6 L of cell culture were harvested by centrifugation (3500 x g) for 20 minutes and resuspended in a buffer containing 20 mM HEPES pH 8.0, 500 mM NaCl, 0.5 mM TCEP, 5 mM imidazole, 1 mM MgCl_2_, cOmplete Protease Inhibitor (Roche), and 10 µg/mL DNase I. Cells were lysed using a Continuous Flow Cell Disruptor (Constant Systems) at 30,000 psi and the lysate was clarified by centrifugation at 48,000 × g for 30 min. at 4 °C and loaded onto a HisTrap HP (Cytiva) column. The column was washed with 100 mL of 20 mM HEPES pH 8.0, 500 mM NaCl, 0.5 mM TCEP and 10 mM imidazole to remove impurities. C-avi-CRBN^midi^ was eluted by a gradient increasing the concentration of imidazole to 150 mM over100 mL. Fractions containing C-avi-CRBN^midi^ were verified using SDS-PAGE and the pooled fractions were diluted 2-fold in a buffer without imidazole to immediately reduce the imidazole concentration to ∼55 mM. 0.3 mg of His-TEV protease was added, and the sample was dialysed in 2 L of 20 mM HEPES pH 8.0, 500 mM NaCl, 0.5 mM TCEP at 4°C overnight. The TEV protease, uncleaved protein, His_6_ tag, and remaining impurities were removed by reverse Ni^2+^ affinity chromatography using HisTrap HP column with C-avi-CRBN^midi^ eluting in buffer containing 15 mM imidazole, 20 mM HEPES pH 8.0, 500 mM NaCl, and 0.5 mM TCEP. 1 mL sample of protein at 100 μM was buffer exchanged using an Amicon Ultra 10 K concentrator (Millipore) to reduce the NaCl concentration to 80 mM for biotinylation with BirA. Final concentrations of 50 mM Bicine pH 8.3, 50 mM magnesium acetate, 2 mM ATP and 150 μM D-biotin were added to the sample along with 160 μg GST-BirA. The biotinylation reaction was incubated at room temperature for 1 hour before a further 160 μg of GST-BirA and 3 μL of 50 μM D-biotin were added and a second incubation implemented. GST-BirA was removed from the sample by a 30-minute incubation at room temperature with 400 μL of glutathione agarose beads. Biotinylated C-avi-CRBN^midi^ was further purified by size-exclusion chromatography (SEC) on a Superdex 75 column (Cytiva) in buffer containing 20 mM HEPES pH 7.5, 500 mM NaCl, and 0.5 mM TCEP. Successful biotinylation was confirmed by a streptavidin gel-shift assay. The protein was concentrated to around 17 μM, flash frozen in liquid nitrogen and stored at -80 °C until further use. All chromatography purification steps were performed using an Äkta Pure (Cytiva) system at 4 °C.

##### CRBN:DDB1^ΔBPB^

Plasmid containing expression cassettes for N- terminally His6 tagged CRBN (2-442) followed by a TEV cleavage site and untagged DDB1^ΔBPB^ (1-395 GNGNSG 706-1140) was constructed from a pFastBac derivative containing His6 tagged CRBN (2-442) and untagged full length DDB1 received from Jana Neuhold (Vienna Biocenter Core Facilities GmbH) via Gibson assembly.

Bacmids were generated by transposition of *E. coli* DH10EMBacY (Geneva Biotech). Baculovirus was produced by transfection of SF9 cells with PEI Max (Sigma Aldrich). For expression, High Five cells at a density of 2 × 10^6^ ml^−1^ were infected with baculovirus. The cells were incubated post infection for 72 h at 27 °C. The cells were collected by centrifugation and stored at -70°C until further use.

Cells were lysed via freeze thaw in 50 mM Tris, 200 mM NaCl, 0.25 mM TCEP, 20 mM Imidazole, pH 8 (buffer A). Recombinant protein was purified by immobilized metal affinity chromatography (Ni Sepharose FF, Cytiva). Affinity resin was washed with buffer A and bound protein was eluted in the same buffer containing 500 mM Imidazole. For anion exchange chromatography using a 5 ml HiTrap Q column (Cytiva), the buffer was exchanged to 50 mM Tris, 130 mM NaCl, 0.25 mM TCEP, pH 8. Protein complex was eluted with a linear gradient to 50 mM Tris, 500 mM NaCl, 0.25 mM TCEP, pH 8 over 20 column volumes.

His tag was removed by incubation with His- tagged TEV protease at 4°C overnight. Reverse affinity was performed using a 5ml HisTrap FF column (Cytiva). Column was washed with buffer A + 20 mM Imidazole. Flow through and wash fraction containing CRBN-DDB1 complex was subjected to size exclusion chromatography using a Superdex 200 16/ 600 column (Cytiva) pre- equilibrated with 50 mM Tris, 200 mM NaCl, 0.25 mM TCEP. Protein was concentrated to 12.4 mg/ ml, flash frozen in liquid Nitrogen and stored at -70°C.

### Protein crystallography

CRBN^midi^ at 3.4 mg/ml (91 uM) in a buffer containing 20 mM HEPES pH 7.5, 500 mM NaCl and 0.5 mM TCEP was mixed with a 4-molar excess of Compound 9 (364 uM, 1.8 % (v/v) DMSO final). The complex was subjected to co-crystallisation across several sparse matrix screens using the sitting drop vapour diffusion method at 20 °C by mixing equal volumes of protein solution and reservoir solution. Initial crystal hits from JCSG+ B4 (Molecular Dimensions) were optimised by a 2D grid screen, and the best diffracting crystals resulted from a reservoir solution containing 0.1 M HEPES pH 7.5, 5.9 % (w/v) PEG 8000 and 10 % ethylene glycol. The crystals were cryoprotected using reservoir solution supplemented with 25 % PEG 400 and flashed cooled in liquid nitrogen.

Diffraction data were collected at SLS X10SA using an EIGER2 16 M detector at 1.00006 Å wavelength. The 2.36 Å dataset was processed using autoPROC^34^, indexed using XDS^35^ and analysed using STARANISO ^36^The structure was solved by molecular replacement in phenix.phaser (version 1.20.1-4487)^37^ using 8RQA as a search model. Phenix.refine (version 1.20.1-4487)^38^ and coot (WinCoot 0.9.4.1)^39^ were used for refinement and model building. Ligand restraints were generated using grade2 (version 1.4.1).^40^ The Ramachandran statistics for the final structure were 94.44 % favoured, 5.24 % allowed and 0.32 % outliers.

### Cryo-electron microscopy sample preparation

Sample preparation was based on Watson *et al*.^5^ CRBN:DDB1^ΔBPB^ was diluted to 50 μM (for Compound 8 and Compound 9) or 34 μM (for Compound 4) and a 5-10-fold molar excess of ligand was added in a final buffer containing 20 mM HEPES pH 7.5, 200 mM NaCl, 0.25 mM TCEP, 1 % DMSO. Samples were diluted 10-fold in DMSO free buffer containing 0.011 % Lauryl Maltose-Neopentyl Glycol (LMNG) immediately before plunge-freezing in liquid ethane. UltrAuFoil R 1.2/1.3 Gold foil on Gold 300 mesh grids (Quantifoil) were glow discharged for 60 s with a current of 35mA under vacuum using a Quorum SC7620 or EasiGlow (PELCO). Grid freezing was carried out using a Vitrobot Mark IV (Thermo Fisher Scientific) with the chamber at 4 °C and 100 % humidity using a blot time of 3s. The grids were pre-treated with CRBN agnostic mutant Ikaros 140-196 Q146A G151N first described by Watson *et al*.,^5^ For CRBN:DDB1^ΔBPB^:Compound 8, 2.4 μL of CRBN agnostic mutant Ikaros was applied to each side of the grid followed by a 60 s wait time before blotting from both sides using Grade 595 Vitrobot filter paper (Ted Pella). 2.4 μL of CRBN:DDB1^ΔBPB^:Compound 8 sample was then applied to each side with a wait time of 0.5 s before blotting from each side and plunging into a liquid ethane pool cooled by liquid nitrogen. For CRBN:DDB1^ΔBPB^:Compound 9, 4 μL of CRBN agnostic mutant Ikaros was applied to one side of the grid and manually blotted from behind after a 60 s wait. 4 μL of CRBN:DDB1^ΔBPB^:Compound 9 sample was then applied to one side, blotted from both sides before plunge freezing.

### Cryo-electron microscopy data acquisition

Cryo-EM data were collected on Krios G4 (for CRBN:DDB1^ΔBPB^:Compound 8) and Glacios (for CRBN:DDB1^ΔBPB^:Compound 9) transmission electron microscopes (Thermo Fisher) operating at 300 keV and 200 keV, respectively. Micrographs were acquired using a Falcon4i direct electron detectors both equipped with Selectris energy filters in counting mode using EPU data collection software (Thermo Fisher) on both the Glacios and Krios. For CRBN:DDB1^ΔBPB^:Compound 8, a total of 6,000 movies were collected at 165,000× magnification with the calibrated pixel size of 0.75 Å per pixel. Images were taken over a defocus range of -0.8 to -1.8 µm, with a total accumulated dose of 50 e^-^Å^2^. For CRBN:DDB1^ΔBPB^:Compound 9, a total of 3,752 movies were collected at 190,000× magnification with the calibrated pixel size of 0.71 Å per pixel. Images were taken over a defocus range of -1.7 to 3.2 µm, with a total accumulated dose of 21.94 e^-^Å^2^. Cryo-EM data collection, refinement and validation statistics can be found in Tables S8-9.

### Cryo-electron microscopy image processing

Data were processed using cryoSPARC (version 4.7.0)^41^ and data processing workflow diagrams can be found in Figures S3 and S5 for CRBN:DDB1^ΔBPB^:Compound 9 and CRBN:DDB1^ΔBPB^:Compound 8, respectively.

#### CRBN:DDB1^ΔBPB^:Compound 9

Movies were motion-corrected with Patch Motion Correction and CTF-corrected using Patch CTF. Poor micrographs were rejected using Manually Curate Exposures based on poor CTF fit and outliers of other metrics. Blob Picker was used on a subset of 100 micrographs and particles were extracted (down sampled 360 to 120 pixel box size) and subjected to 2D classification. Promising classes were used as templates for template picking on all accepted micrographs (2,330). 1,283,207 particles were extracted with (down sampled 360 to 128 pixel box size) 2D classification. 12 promising 2D classes were manually selected and run through the Rebalance 2D job resulting in 219,734 particles for Ab-Initio model generation with 5 classes followed by heterogeneous refinement. A class showing good density for both CRBN and DDB1^ΔBPB^ was used to generate 50 2D templates for template picking. 1,585,848 particles were extracted with 360 pixel box size and down sampled to 128 pixels and filtered through iterative rounds of Ab-Initio model generation and Heterogeneous Refinement followed by 3D Classification to yield final subsets of 86,485 and 56,510 particles with CRBN in the open or closed conformation, respectively. The CRBN open particle set was subjected to Global CTF Refinement fitting tilt, trefoil and anisotropic magnification, Local CTF Refinement, Reference-Based Motion Correction^42^ and Non-Uniform Refinement to yield a consensus refinement with a global resolution of 3.33 Å (accession code EMD-55059). The CRBN closed particle set was subjected to Global CTF Refinement fitting tilt, trefoil and anisotropic magnification, Reference-Based Motion Correction^42^ and Non-Uniform Refinement to yield a consensus refinement with a global resolution of 3.66 Å (accession code EMD-55057). Local masks were created for the TBD or LON-TBD in ChimeraX for open and closed CRBN, respectively. Local refinement for TBD or LON-TBD was performed with default settings other than using pose/shift Gaussian prior during alignment with a rotation search extent of 6° (standard deviation of prior over rotation: 6°) and shift search extent of 6 Å (standard deviation of prior over shifts: 2 Å), and re-center rotations and shifts each iteration turned on. The resolution of the locally refined TBD map for the open conformation (accession code EMD-55060) and LON-TBD map for the closed conformation (accession code EMD-55058) was 4.23 Å and 4.07 Å, respectively. Gold-standard FSC, cFSC, angular distribution, and posterior position directional distribution plots for the consensus and locally refined maps of both the open and closed conformation can be found in Figure S4.

#### CRBN:DDB1^ΔBPB^:Compound 8

Movies were motion-corrected with Patch Motion Correction and mrc micrograph files were transferred between sites and reimported as micrographs into cryoSPARC. Micrographs were CTF-corrected using Patch CTF, and poor micrographs were rejected using Manually Curate Exposures based on poor CTF fit and outliers of other metrics. Blob Picker was used on a subset of 500 micrographs and particles were extracted (down sampled 360 to 120 pixel box size) and subjected to 2D classification. Promising classes were used as templates for template picking on all accepted micrographs (4,735). 2,575,103 particles were extracted with 360 pixel box size and down sampled to 120 pixels for Ab-Initio model generation with 8 classes followed by heterogeneous refinement. A single class containing 766,006 particles was taken forward and subjected to Homogeneous Refinement at the full box size of 360 pixels. The particles were further classified using 3D Classification (4 classes). 3 of the 4 classes (573,499 particles) displayed map density for the TBD domain and were pooled for Non-Uniform Refinement.^43^ Subsequent Global CTF Refinement fitting tilt, trefoil, spherical aberration, tetrafoil and anisotropic magnification followed by Local CTF Refinement and a final Non-Uniform Refinement yielded a consensus map (accession code EMD-54624) at 2.22 Å global resolution. A mask was created for local refinement of the TBD in ChimeraX. Local refinement of the TBD was performed with default settings other than using pose/shift Gaussian prior during alignment with a rotation search extent of 9° (standard deviation of prior over rotation: 3°) and shift search extent of 6 Å (standard deviation of prior over shifts: 2 Å), and re-center rotations and shifts each iteration turned on. The resolution of the locally refined map was 3.44 Å (accession code EMD-54644). Gold-standard FSC, cFSC, angular distribution, and posterior position directional distribution plots for the consensus and locally refined maps can be found in Figure S6.

### Cryo-electron microscopy model building

#### CRBN:DDB1^ΔBPB^:Compound 9 open conformation

To combine consensus and focused maps in ChimeraX, the volume multiply command was used on the locally refined map using the local mask used in refinement around the TBD. The vop maximum command was then used to create a composite map (accession code EMD-55259) of the consensus and mask multiplied TBD map. A starting model was created using the CRBN:DDB1^ΔBPB^:Compound 8 structure with the TBD region was replaced by the TBD from the CRBN^midi^:Compound 9 crystal structure. Ligand restraints previously generated for the CRBN^midi^:Compound 9 were used. Due to low signal for the CRBN LON domain and consistent with previous open apo structure of CRBN^5^ most of the domain was modelled with side chains truncated. The model was refined with iterative rounds of manual building in Coot and refinement with Phenix using phenix.real_space_refine.

#### CRBN:DDB1^ΔBPB^:Compound 9 closed conformation

Using the mask for local refinement of the LON-TBD module of CRBN, a composite map (accession code EMD-55258) was generated by the same method as the CRBN:DDB1^ΔBPB^:Compound 9 open conformation. The initial model was created using the CRBN:DDB1^ΔBPB^:Compound 8 structure, replacing the corresponding regions of CRBN with the high-resolution X-ray model of CRBN^midi^:Compound 9. The model was refined in the same manner as the open conformation structure.

#### CRBN:DDB1^ΔBPB^:Compound 8

The vop maximum command in UCSF ChimeraX ^44^(version 1.10.1) was used to combine the consensus and focused TBD maps into a composite map (accession code EMD-55257). A cryo-EM structure of apo CRBN:DDB1 in the open conformation (PDB code: 8CVP^5^ was used as an initial model. Grade2 (version 1.4.1)^40^ was used to generate ligand restraints and the model refinement was performed with iterative rounds of manual building in Coot (WinCoot 0.9.8.95, ^39^) and refinement with Phenix (version 1.20.1-4487,^45^) using phenix.real_space_refine. Due to weak signal for the CRBN LON domain and consistent with previous open apo structure of CRBN^5^ most of the domain was modelled with side chains truncated. To improve the compound pose and geometry, the model was prepared in Maestro (Schrödinger Release 2024-4, Schrödinger, LLC, New York, NY, 2025.) and ligand minimization was performed. The ligand was output as a mol2 file from Maestro and restraints were generated from the input geometry and used in the final refinement of the model.

### SAXS

SAXS data were collected in SEC-SAXS configuration at beamline B21 at the Diamond Light Source. ^46^ CRBN^midi^:binder complexes were formed by the addition of 400 μM of binder to 2.4-3.4 mg/mL of CRBN^midi^ in a final buffer consisting of 20 mM HEPES pH 7.5, 500 mM NaCl, 0.5 mM TCEP and 4 % DMSO. Samples were run on a Superdex S200 3.2/300 column (Cytiva) in SAXS buffer with 0.075 mL/min. flowrate at 298 K coupled online to the SAXS beamline. SAXS data were acquired using a 3 s exposure at 12.4 kV and 3.6 m detector (Eiger 4 M) distance across a q-range of 4.5 × 10^-3^ - 3.4 × 10^-1^ Å^-1^. Frames from SEC-SAXS that corresponded to the main protein peak, were merged using Chromixs^47^ and analysed using the ATSAS package^48^ Radii of gyration and molecular mass were extrapolated from the Guinier plot using Primus. Pairwise-distance distributions were calculated using GNOM.^49^ The dimensionless Kratky plot^50^ was obtained by manually plotting (qR_g_)^2^I(q)/I_0_ against qR_g_.

### Structural Analysis

The super command in PyMOL (version 2.4) was used to calculate the RMSD values for structural alignment. Structure figures were generated using PyMOL (version 2.4) or UCSF ChimeraX (version 1.10.1).

### Surface Plasmon Resonance

SPR measurements with CRBN^midi^ were carried out on a Biacore^TM^ 8 K (Cytiva, Biacore Insight Control Software version 5.0.18.22405). Biotinylated C-avi-CRBN^midi^ was immobilised on a SAD200M chip (XanTec) at 20 °C. The chip surface was prepared for immobilisation by flowing a mixture of 1 M NaCl and 50 mM NaOH followed by buffer, and then 6 M Guanidine HCl also followed by buffer. In each step the contact time was 600 s and the flow rate was 10 μL/min. Biotinylated C-avi-CRBN^midi^ was diluted to 150 nM and flowed over flow cell 2 at a rate of 10 µL/min for 500 s to capture a final bound response of ∼6000 RUs. The buffer for immobilisation and all affinity measurements was 50 mM Tris pH 8.0, 150 mM NaCl, 0.25 mM TCEP, 0.005% (v/v) Tween-20 and 2% (v/v) DMSO.

Steady state affinity measurements were derived from a multi-cycle affinity measurement, with a 5-point 3-fold dilution series of analyte. The top concentration for each binder varied dependent on the *K*_D_. Each concentration point was flowed independently at 50 µL/min. with an association time of 150 s followed by 400 s dissociation. All measurements were carried out at 20 °C.

Data was analysed using Biacore^TM^ Insight Evaluation Software (version 5.0.18.22102). Raw sensorgrams were solvent corrected, followed by reference and blank subtraction. To calculate *K*_D_ values, data was globally fitted to a steady state affinity model.

SPR experiments with CRBN^TBD^ were performed with a Bruker SPR32 instrument. N-terminal His-tag was cleaved by TEV-protease and revealed N-avi-CRBN^TBD^ which was biotinylated and immobilized on a XanTec SAD-200M chip. The running buffer for immobilisation and affinity determinations was 20mM TRIS pH 7,5, 150mM NaCl, 0,005% Tween-20, 1mM DTT using a flow rate of 5µL/min. Additional 2% (v/v) DMSO were present for compound interaction experiments. Conditioning the sensor chip was done by injecting 1M NaCl/40mM NaOH over for three times 60s, followed by injection of running buffer for 2×60s. All spots of each flow cell were treated equally. The protein was diluted to around 0,055 mg/ml using running-buffer for immobilization and injected for 180s over spots B,C and D of each flow cell. Lastly, running buffer was injected twice for 60s over all spots of each flow cell.

Steady state K_D_ measurements were carried out as multi cycle kinetic experiments with 11 increasing concentrations using an association time of 60s with a flow rate of 30µl/min. The dissociation time was adjusted to the off rate of individual compounds. For data analysis sensorgrams from reference surfaces and blank injections were subtracted from the raw data prior to data analysis, using SPR32 Analyser R2. The steady-state-affinity model was used to calculate the K_D_, since all compounds showed fast binding kinetics.

For BRD4 morpholinone PROTAC analysis biotinylated C-avi-CRBN^midi^ was diluted to 12.5 nM and flowed over flow cell 2 at a rate of 10 µL/min for 500 s to capture a final bound response of ∼ 400 RUs. The buffer for immobilisation and all kinetic measurements was 50 mM Tris pH 8.0, 150 mM NaCl, 0.25 mM TCEP, 0.005% (v/v) Tween-20 and 2% (v/v) DMSO.

Kinetic measurements were derived from a multi-cycle affinity measurement, with a 5-point 3-fold dilution series of analyte (500 nM-6.17nM for morpholinone PROTACS, 5000 nM-6.17 nM for CTF1297). For ternary measurements the PROTAC was saturated with 2 μM BRD4 BD2. Each concentration point was flowed independently at 40 µL/min. with an association time of 180 s followed by 1200 s dissociation. All measurements were carried out at 20 °C.

Data was analysed using Biacore^TM^ Insight Evaluation Software (version 4.0.8.20368). Raw sensorgrams were solvent corrected, followed by reference and blank subtraction. To calculate *K*_D_ values, data were globally fitted to a 1:1 binding kinetic model.

### Differential Scanning Fluorimetry

Samples were prepared in a total volume of 15 µL by mixing 10 µM of CRBN^midi^ with 100 µM of compound when applicable in a final buffer containing 50 mM HEPES pH 7.5, 200 mM NaCl, 0.5 mM TCEP and 4% (v/v) DMSO. Samples were loaded into Prometheus standard capillaries (NanoTemper Technologies) and thermal unfolding experiments were carried out by increasing the temperature from 20 °C to 95 °C at a rate of 1 °C/min. using a Prometheus Panta (NanoTemper Technologies). Protein aggregation was assessed, and turbidity values were calculated by the PR.Panta control software (NanoTemper Technologies). First derivative curves were plotted using Microsoft Excel.

Thermal melting curves for CRBN^TBD^ was carried out employing a Bio-Rad CFX384 instrument using white 384-Well PCR PP plates (Brand) sealed by Microseal®’B’ Adhesive Seals (Bio-Rad). The protein was pre-mixed with SYPRO orange DMSO stock solution (SIGMA, 5000x) yielding a CRBN^TBD^ concentration of 50µM and a 25fold excess of the dye. Adding 2µL of the protein dye mix to 8ul of the pre-diluted compound solution revealed final concentrations of 10µM protein and 500µM compound as well as 2% (v/v) DMSO. Plates were centrifuged for 2 min at 1000g and subjected to a temperature ramp starting from 15°C to 95°C at a rate of 1 °C/min and fluorescence measurements every 0.5°C. Thermal stabilisation of compounds were assessed by the difference of the melting temperature (T_m_) of CRBN^TBD^ in the presence of the compound and the average of multiple DMSO negative controls. Data analysed by the Bio-Rad CFX Manager Software package and exported to Excel for delta T_m_ calculation.

### Isothermal Titration Calorimetry

Experiments were carried out on an ITC200 instrument (Malvern), using MicroCal ITC200 (version 1.26.1) software, in buffer containing 20 mM HEPES pH 7.5, 200 mM NaCl, 0.25 mM TCEP, 2% (v/v) DMSO at 298 K stirring the sample at 600 rpm. An initial 0.4 µL injection (discarded during data analysis) was followed by 19 × 2 µL injections with 180 s spacing between injections. CRBN binders (400 µM) was directly titrated into CRBN:DDB1 (30 µM). For a control titration, 400 µM of CRBN binders were titrated into buffer. The data were fitted using a one-set-of-site binding model to obtain dissociation constants, binding enthalpy (ΔH), and stoichiometry (N) using MicroCal PEAQ-ITC Analysis Software (version 1.1.0.1262, Malvern).

### NMR

The protein NMR detected fragment screen was carried out on a Bruker Avance Neo 600 MHz equipped with a 1.7mm TCI cryogenic probe at 298 K. Samples were prepared just in time before data acquisition with an in-house modified Tecan Fluent pipetting robot and automatically transferred into the respective sample tube, which was then transported to the magnet via an adapted Bruker SampleMail sample changer. The in house generic fragment library of 8340 molecules was screened in preformed mixtures of ten at a final concentration of 500 µM of each compound (2.5 % (v,v) d6DMSO) and 30 µM ^15^N labeled CRBN^TBD^ in 50mM Tris-d11, 200mM NaCl, 0.5mM TCEP pH 7.5 and 10% D2O. Mixtures experiencing significant chemical shift perturbation or line broadening were deconvoluted by measuring individual compounds under the above mentioned experimental conditions.

### Cell culture and Stable Cell line Generation

All cell lines were maintained in a humidified incubator at 37 °C and 5% CO_2_, authenticated by STR profiling and regularly tested for mycoplasma contamination. DLD-1 WT and DLD-1 CRBN KO lines were obtained from MRC PPU, University of Dundee. HEK293 were obtained from LGC (CRL-1573). All cells were grown in Dulbecco’s Modified Eagles Medium (DMEM, 11965092, Thermo) containing 10% (v/v) Foetal Bovine Serum (FBS, A5256801, Thermo).

For creation of FLAG-CRBN expressing DLD-1 lines, viruses were initially generated in HEK 293 FT cells followed by subsequent viral transduction of DLD-1 CRBN KO cells. To generate virus, HEK 293 FT cells were grown to 50-60% confluency in one 10 mm dish per virus. The following day, a mixture containing 3 µg pBABED plasmid, 3.8 ug GAG/POL, 2.2 µg VSVG and 24 µl polyethyleneimine (PEI; Polysciences) was diluted into 500 µl Opti-MEM reduced serum medium (Gibco) and incubated for 30 min at room temperature before adding dropwise onto cells. 24h after transfection, the transfection-containing medium was discarded, and 10 ml fresh medium was added. 24h later, virus containing medium was harvested and filtered using a 0.45 µm filter before storage.

For viral transduction, DLD-1 CRBN KO cells were plated at a density of 0.75 x 10^6^ cells per well into a 6-well plate, with 3 wells per condition. The following day, 2 ml virus containing polybrene (8 µg/ml) was added to one well, 1 ml virus containing polybrene (8 µg/ml) and 1 ml fresh medium added to the second well and 2 ml fresh medium only added to the third. After 24 h cells were selected for positive viral uptake by replacing media with fresh medium containing 2 µg/ml puromycin (Invivogen; #ant-pr-1) until all the cells in the un-transduced control well were dead.

### NanoBRET™ Target Engagement Assay

Binding affinity and cell permeability of CRBN-based PROTACs were assessed using the NanoBRET™ Target Engagement (TE) assay (Promega) in intact HEK293 cells stably expressing NanoLuc-CRBN fusion protein. The NanoBRET™ (bioluminescence resonance energy transfer) TE Assay is a proximity-based assay measuring the energy transfer from a donor (luciferase) to an acceptor (fluorescent protein or probe). In this setup, the basis of the NanoBRET™ TE Assay is the competitive displacement of a fluorescent CRBN-Tracer reversibly bound to a CRBN-NanoLuc fusion protein (bioluminescent donor) in cells. If the tracer is bound to CRBN and therefore in close proximity to the NanoLuc® luciferase, energy transfer generating the NanoBRET™ signal occurs. When the tested compounds bind to CRBN, they displace the tracer, resulting in loss of the NanoBRET™ luminescence signal. HEK293 NanoLuc-CRBN cells were cultured in DMEM supplemented with 10% FBS and 800 µg/mL Geneticin, maintained at 37°C and 5% CO₂. Cells were passaged every 3–4 days and used up to passage 25. For the assay, 10,000 cells per well in 40 µl were seeded in 384-well white polystyrene plates (Corning, PN: 3574) in Opti-MEM. After 24 hours, cells were incubated with 350 nM NanoBRET™ CRBN tracer (Promega, PN: N2912) for 15 seconds on a microplate shaker. PROTACs were added via Echo® 555 Liquid Handler in a 1:3 serial dilution (top concentration: 10 µM) and incubated for 2 hours at 37°C. Following compound incubation, NanoBRET™ Nano-Glo® Substrate and Extracellular NanoLuc® Inhibitor were added (final dilution: 1:166 and 1:500, respectively). Plates were equilibrated to room temperature for 15 minutes, shaken briefly, and luminescence was measured using an EnVision® 2105 plate reader (PerkinElmer) at donor (460 nm) and acceptor (615 nm) wavelengths. Negative controls (0.5% DMSO) were included on each plate. Edge wells were excluded from compound treatment to minimize variability.

### TR-FRET Assay to assess competitive binding to CRBN+DDB1 Complex

A time-resolved fluorescence resonance energy transfer (TR-FRET) assay was developed to identify compounds that competitively inhibit the binding of Cy5-labeled lenalidomide to the CRBN+DDB1 complex. The assay utilizes a 6xHis-tagged CRBN+DDB1 protein labeled with a europium-conjugated anti-His antibody (fluorescence donor) and Cy5-labeled lenalidomide (fluorescence acceptor). Signal intensity correlates with the proximity of donor and acceptor, reflecting binding interactions. Assay buffer consisted of 20 mM HEPES (pH 7.3), 150 mM NaCl, and 0.005% Tween-20 in UltraPure water. The CRBN+DDB1 complex was used at 60 nM. Europium-labeled anti-6xHis antibody (Revvity, AD0400) was added at 3 nM. Cy5-labeled lenalidomide was used at 30 nM, prediluted 1:100 in DMSO. Assays were performed in 384-well white Proxiplates (PerkinElmer, PN: 6008280) with a total reaction volume of 15 µL. Dose-response curves were generated using 10 mM DMSO stock solutions dispensed via Echo® 55x/65x Liquid Handler (Beckman Coulter). Eleven concentrations were tested in 1:5 serial dilutions in technical duplicates. Cy5-labeled lenalidomide was added to columns 1–23; column 24 served as a blank control. Plates were processed in a darkened room (<100 Lux) using a fully automated robotic system. After addition of CRBN/Eu-Mix, plates were mixed for 20 seconds at 1750 rpm and incubated for 42 minutes at room temperature. TR-FRET signals were measured using an EnVision® 210x multimode plate reader (PerkinElmer) and analyzed using an inhouse LIMS software. Negative controls included reactions with DMSO and Cy5-labeled binder; positive controls excluded the Cy5-labeled binder. Reference compounds were tracked in MegaLab and used to monitor assay stability.

### Western Blot

Cells were seeded into 6 or 12-well plates at a density of 0.3×10^6^ cells/ml 24h prior to compound treatment. The following day, cells were treated with DMSO or compounds and incubated for the indicated time. Following incubation, cells were washed twice with PBS and lysed in lysis buffer (1% Triton X-100, 150 mM NaCl, 1 mM EDTA, 50 mM Tris pH 7.4, protease inhibitor cocktail (Roche) and phosphatase inhibitor tablets (Merck; #4906845001). Lysates were clarified by centrifugation at 17,000 x g for 15 min at 4°C and supernatant stored for further analysis.

Protein concentration of cell lysates was determined by BCA assay (Pierce) and the absorbance at 562 nm measured using a Pherastar (BMG Labtech). Cell extracts were denatured in lithium dodecyl sulfate (LDS) (NuPAGE) containing 25 nM DTT followed by incubation for 5 min at 95 °C to denature proteins and reduce disulphide bonds. Samples were separated by SDS-PAGE using 20 μg of protein per well of NuPAGE Novex 4–12% BIS-TRIS gels (Invitrogen) and transferred to 0.2 μm pore nitrocellulose membrane (Amersham) by wet transfer (90V for 90min).

Membranes were probed with primary antibodies targeting BRD4 ( # 1:1000), CRBN (CST; #71810, 1:1000). Rhodamine-Tubulin (BioRad; #12004165). Rhodamine-GAPDH (BioRad; #12004167). Secondary antibodies coupled to fluorophores were used; IRDye 800 donkey anti-rabbit secondary (LI-COR; #926-32213). Western blots were developed using a ChemiDoc MP imaging system (Bio-Rad) and quantified using ImageLab (Bio-Rad) or Image Studio (LI-COR) with analysis using GraphPad Prism (version 10.6.0). Original western blots are provided as supplementary material.

### Plasmids and Constructs

pCMV VSV-G (# RV-110) and pCMV VSV-G retrovirus envelope plasmids (# RV-110) were purchased from Cell BioLabs. pBABED FLAG-CRBN (#DU54685), pBABED FLAG-CRBN L60A (#DU79417), pBABED FLAG-CRBN H378N (#DU79418), pBABED FLAG-CRBN Q100A (#DU79419), pBABED FLAG-CRBN H378A (#DU79427) and pBABED FLAG-CRBN L60A/H378A (#DU79428) were cloned and purchased from MRC PPU Reagents and Services, University of Dundee.

