## Supporting Information for "Tuning the open-close equilibrium of Cereblon with small molecules influences protein degradation"

Table S3. Measurement of different CRBN classes by DSF with CRBN$^{midi}$. Turbidity first derivative of thermal denaturation for CRBN$^{midi}$ in the absence or presence of binders. .. 13

### Chemical Synthesis & Reagents

#### General Information

Commercially available dry solvents were obtained from Sigma Aldrich. All reagents, unless otherwise noted, were commercially available and purchased from Sigma Aldrich, Combi Blocks, ABCR, Fluorochem, Activate or Enamine, at least 95% pure and used without further purification. All reactions were carried out under a nitrogen atmosphere. Normal phase TLC was carried out on pre-coated silica plates (Kieselgel 60 F254, BDH) with visualization via UV light (UV 254 and/or 365 nm) and/or basic potassium permanganate solution. Isolute® phase separator columns from Biotage were used. Flash column chromatography was performed using either a Teledyne Isco Combiflash Rf or Rf200i, or a Biotage Isolera One with prepacked Redisep RF normal phase disposable columns. Reverse phase chromatography was carried out using Biotage SNAP-C18 columns or RediSep Rf Reversed Phase C18 Columns. Strong cation exchange (SCX) chromatography was carried out using Biotage Isolute SCX-2 columns. NMR Spectra were recorded on Bruker 400 MHz or 500 MHz spectrometers as specified. Chemical shifts are quoted in ppm and referenced to the residual solvent signals: $^{1}$H NMR δ (ppm) = 7.26 ($CDCl_3$-d), 5.32 ($CD_2Cl_2$) or 3.31 (MeOD-$d_4$). Signal splitting patterns are described as singlet (s), doublet (d), triplet (t), quartet (q), quintet (quin.), multiplet (m), broad (br) or a combination thereof. Coupling constants (J) are measured in Hertz (Hz). Diastereomeric ratios (dr) were calculated using the ratios of NMR integrals. Chiral SFC was carried out by Reach Separations Ltd., using a Sepiatec SFC systems equipped with a Lux A1 column (21.2 mm x 250 mm, 5 um particle size). Samples were eluted with an isocratic gradient of 40:60 EtOH:$CO_2$ (0.2% v/v $NH_3$) over 10 min at a flow rate of 50 mL/min with an oven temperature of 40 °C. Diastereomeric mixture was dissolved in EtOH at 6 mg/mL and injected 16 times at 500 uL.

##### *Abbreviations*

aq. for aqueous, Boc for *N*-*tert*butyloxycarbonyl, DCM for dichloromethane, DMA for *N*,*N*-dimethylacetamide, DMF for *N*,*N*-dimethylformamide, DIPEA for N,N-diisopropylethylamine, DMSO for dimethylsulfoxide, HATU for 1-[bis(dimethylamino)methylene]-1*H*-1,2,3-triazolo[4,5-b]pyridinium 3-oxide hexafluorophosphate, TBAF for tetra-*N*-butylammonium fluoride

#### Analytical MS Methods and instrumentation

##### *LCMS*

Liquid chromatography-mass spectrometry (LCMS) was carried out on a Shimadzu HPLC/MS 2020 equipped with a Hypersil Gold column (1.9 µm particle size, 50 × 2.1 mm), photodiode array detector and ESI detector. Samples were eluted with either a 3 min or 5 min gradient of 5–95% acetonitrile:water containing 0.1% formic acid at a flow rate of 0.7 mL/min.

##### *HPLC*

Preparative HPLC was performed on a Waters Prep 150 LC system with a Waters XBridge C18 column (100 mm x 19 mm; 5 µm particle size) and a gradient of 5% to 95% acetonitrile in water over 20 minutes and a flow rate of 25 mL/min, with ammonia in the aqueous phase.

#### Experimental Procedures

Compound **1** is commercially available (CAS: 443955-72-4) and was purchased from Enamine.

Synthesis of compounds **9**, **10**, **11** and **12** were conducted according to literature references. [1–4]

**Synthesis of 7-(4-Methylpiperazine-1-carbonyl)-1H,2H,3H-pyrido[2,3-b][1,4]oxazin-2-one (compound 2)**

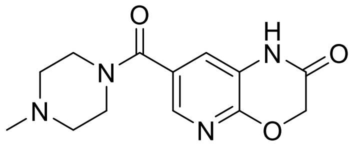

To a solution of 2-oxo-1H,2H,3H-pyrido[2,3-b][1,4]oxazine-7-carboxylic acid (200 µmol, 39.6 mg) in dimethylacetamide (1 mL), N,N-diisopropylethylamine (1160 µmol, 0.151 g, 0.2 mL) was added. The mixture was stirred briefly to ensure dissolution. Subsequently, HATU (210 µmol, 80 mg) was added, and the reaction mixture was stirred briefly to activate the acid. Once the activation was complete, 1-methylpiperazine (300 µmol, 0.031 g, 0.034 mL) was added to the reaction mixture. The resulting solution was stirred at room temperature for 3.5 hours. The progress of the reaction was monitored by LCMS, which confirmed complete conversion of the starting material. The reaction mixture was diluted with acetonitrile-water and subjected to purification using preparative HPLC. The purified product, 7-(4-methylpiperazine-1-carbonyl)-1H,2H,3H-pyrido[2,3-b][1,4]oxazin-2-one, was obtained as a solid (18.1 mg, 33% yield).

MS (ESI+): m/z calcd. for $[M+H^+]$: 277.12, found: 277.19.

$^1$H NMR (400 MHz, DMSO-d6) δ ppm: 10.91 (s, 1H), 7.83 (d, $J$ = 2.0 Hz, 1H), 7.22 (d, $J$ = 2.0 Hz, 1H), 4.83 (s, 2H), 3.36 - 3.78 (m, 4H), 2.26 - 2.43 (m, 4H), 2.21 (s, 3H).

**Synthesis of 7-(1-methyl-1H-1,3-benzodiazol-2-yl)-1H,2H,3H-pyrido[2,3-b][1,4]oxazin-2-one (compound 3)**

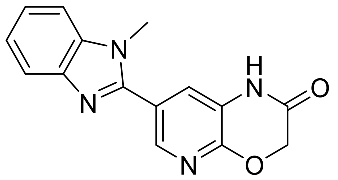

To a solution of 2-oxo-1H,2H,3H-pyrido[2,3-b][1,4]oxazine-7-carboxylic acid (100 µmol, 19.8 mg) in dimethylformamide (2 mL), N,N-diisopropylethylamine (0.3 mmol, 51.55 µL) was added at room temperature. The mixture was stirred briefly to ensure dissolution. Subsequently, HATU (0.11 mmol, 41.8 mg) was added, and the reaction mixture was stirred for 10 minutes to activate the acid. N-methyl-ortho-phenylenediamine (0.1 mmol, 12.2 mg) was then added to the reaction mixture. The resulting solution was stirred at room temperature for 1 hour. The progress of the reaction was monitored by LCMS, which confirmed the formation of the intermediate product. The reaction mixture was filtered through basic alumina and washed with a DMF/methanol mixture (9:1). The filtrate was concentrated under reduced pressure to yield a residue which was treated with acetic acid (2 mL) and refluxed. LCMS analysis confirmed the formation of the desired product. The crude product was concentrated under reduced pressure, dissolved in dimethylformamide, and purified using reverse-phase liquid chromatography on a C18 column with an acetonitrile-water-trifluoroacetic acid (ACN/H2O/TFA) mobile phase at 50°C. The collected fractions were freeze-dried to yield the purified product as a yellow solid (28.9 mg, 73% yield).

MS (ESI+): m/z calcd. for [M+H$^+$]: 281.10, found: 281.19.

$^1$H NMR (400 MHz, DMSO-d6) δ ppm: 11.16 (s, 1H), 8.32 (d, *J* = 2.15 Hz, 1H), 7.85 (d, *J* = 7.86 Hz, 1H), 7.79 (d, *J* = 7.35 Hz, 1H), 7.70 (d, *J* = 2.15 Hz, 1H), 7.43 - 7.56 (m, 2H), 4.94 (s, 2H), 1.96 (s, 3H).

**Synthesis of 7-(1-cyclohexyl-1H-1,3-benzodiazol-2-yl)-1H,2H,3H-pyrido[2,3-b][1,4]oxazin-2-one (compound 4)**

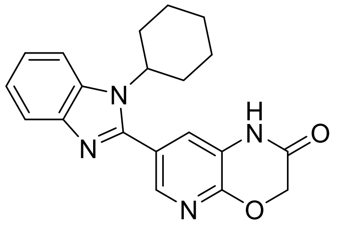

To a solution of 2-oxo-1H,2H,3H-pyrido[2,3-b][1,4]oxazine-7-carboxylic acid (100 µmol, 19.8 mg) in dimethylformamide (2 mL) at room temperature, N,N-diisopropylethylamine (0.3 mmol, 52 µL) was added. The mixture was stirred briefly to ensure dissolution. Subsequently, HATU (0.11 mmol, 41.8 mg) was added, and the reaction mixture was stirred for 10 minutes. N1-cyclohexylbenzene-1,2-diamine (0.1 mmol, 19.03 mg) was then added to the reaction mixture, and the resulting solution was stirred at room temperature for 1 hour. The progress of the reaction was monitored by LCMS, which confirmed the formation of the intermediate product. The reaction mixture was filtered through basic alumina and washed with a DMF/methanol mixture (9:1). The filtrate was concentrated under reduced pressure to yield crude residue. To induce cyclization, the residue was treated with acetic acid (2 mL) and refluxed. LCMS analysis confirmed the formation of the desired product. The crude product was concentrated under reduced pressure, dissolved in dimethylformamide, and purified using reverse-phase liquid chromatography on a C18 column with an acetonitrile-water-trifluoroacetic acid (ACN/H2O/TFA) mobile phase at 50°C. The purified product was obtained as a beige amorphous solid (30.4 mg, 87% yield).

MS (ESI+): m/z calcd. for [M+H$^+$]: 349.16, found: 349.21.

$^1$H NMR (400 MHz, DMSO-d6) δ ppm: 11.18 (s, 1H), 8.16 (s, 1H), 8.11 (dd, *J* = 5.7, 3.0 Hz, 1H), 7.80 (dd, *J* = 6.0, 3.1 Hz, 1H), 7.57 (s, 1H), 7.47 (dd, *J* = 6.0, 3.0 Hz, 2H), 5.14 (s, 1H), 4.95 (s, 3H), 4.22-4.45 (m, 2H), 2.18-2.43 (m, 4H), 2.00 (d, *J* = 11.3 Hz, 2H), 1.86 (d, *J* = 11.2 Hz, 3H), 1.65 (s, 1H), 1.30-1.55 (m, 4H).

**Synthesis of 7-Pyridazin-3-yl-1,7-diazaspiro[4.4]nonan-2-one (compound 6)**

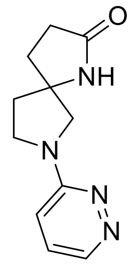

To a 5 mL microwave vial, 3-bromopyridazine (1.0 eq, 0.06 g, 0.378 mmol) and 1,7-diazaspiro[4.4]nonan-2-one (1.2 eq, 0.063 g, 0.452 mmol) were dissolved in N-methylpyrrolidone (2 mL) at room temperature. The resulting solution was degassed with nitrogen. DIPEA (0.415 eq, 0.0106 mL, 0.157 mmol) and potassium iodide (0.100 eq, 0.006 g, 0.0378 mmol) were then added. The reaction mixture was stirred at 170°C for 1.5 hours. The progress of the reaction was monitored by TLC and LCMS, which confirmed the complete consumption of the starting material and the formation of the desired product. The reaction mixture was diluted with ice water (20 mL) and extracted with ethyl acetate (2 × 10 mL). The aqueous layer was lyophilized to yield a crude material which was purified using preparative HPLC to yield the desired product (60 mg, 22%).

MS (ESI+): m/z calcd. for $[M+H^+]$: 219.12, found: 219.16.

$^1H$ NMR (400 MHz, DMSO-d6) δ ppm: 2.06 (m, 4H), 2.29 (m, 2H), 3.43 (m, 1H), 3.51 (m, 2H), 3.58 (m, 1H), 6.83 (dd, J = 9.2, 1.2 Hz, 1H), 7.34 (dd, J = 9.2, 4.4 Hz, 1H), 8.12 (s, 1H), 8.48 (dd, J = 4.4, 1.2 Hz, 1H).

**Synthesis of 2-Pyridazin-3-yl-2,5-diazaspiro[3.4]octan-6-one (compound 7)**

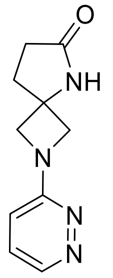

In a 5 mL microwave vial, 3-bromopyridazine (1.0 eq, 0.030 g, 0.189 mmol) and 2,5-diazaspiro[3.4]octan-6-one (1.5 eq, 0.034 g, 0.283 mmol) were dissolved in N-methylpyrrolidone (2 mL) at room temperature. DIPEA (2.00 eq, 0.065 mL, 0.378 mmol) and potassium iodide (0.100 eq, 0.003 g, 0.0189 mmol) were then added. The reaction mixture was stirred at 170°C for 1.5 hours. The progress of the reaction was monitored by TLC and LCMS, which confirmed the formation of the desired product, although the starting material was not completely consumed. The reaction mixture was diluted with ice water (20 mL) and extracted with ethyl acetate (2 × 10 mL). The aqueous layer was lyophilized to yield a crude compound (0.03 g). The crude product was purified using preparative HPLC to give the final product (30 mg, 70%).

MS (ESI+): m/z calcd. for $[M+H^+]$: 205.10, found: 205.15.

$^1$H NMR (400 MHz, DMSO-d6) δ ppm: 2.35 (m, 4H), 4.08 (dd, $J$ = 31.6, 8.8 Hz, 4H), 6.83 (dd, $J$ = 9.0, 1.6 Hz, 1H), 7.38 (dd, $J$ = 9.0, 4.8 Hz, 1H), 8.28 (s, 1H), 8.57 (dd, $J$ = 4.6, 1.6 Hz, 1H).

**Synthesis of 2-(5-chloro-1-(piperidin-4-yl)-1H-indol-4-yl)-2,5-diazaspiro[3.4]octan-6-one (compound 8)**

4-(4-bromo-5-chloro-indol-1-yl)piperidine-1-carboxylate

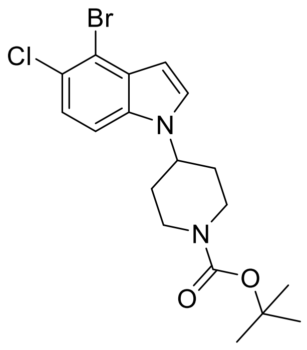

4-bromo-5-chloro-1H-indole (1 eq, 450 mg, 1.95 mmol), tert-butyl 4-methylsulfonyloxypiperidine-1-carboxylate (2 eq, 1091 mg, 3.90 mmol) and $Cs_2CO_3$ (3 eq, 1908 mg, 5.86 mmol) were dissolved in MeCN (15mL) and stirred at reflux under $N_2$ atmosphere for 18 h. A further 500 mg of mesylate was added and the reaction was stirred for 6 h, before the addition of another 500 mg. The reaction mixture was then stirred overnight. The solvent was removed from the reaction mixture and the crude material was taken up in EtOAc (30 mL) and water (30 mL). The organic layer was washed with water (3 x 30 ml) and then brine (30 ml) before being dried over anhydrous $Na_2SO_4$. The crude material was purified by flash reverse phase chromatography, elution 10 to 95% ACN in water + 0.1% formic acid, to afford the product tert-butyl 4-(4-bromo-5-chloro-indol-1-yl)piperidine-1-carboxylate (375 mg, 44%) as a colourless oil.

LCMS: m/z calcd. for [M+H$^+$] 413.7, found: 412.9; RT = 2.24 min.

$^1$H NMR (500 MHz, $CDCl_3$) δ ppm: 7.26 (d, $J$ = 4.2 Hz, 2H), 7.25 (d, $J$ = 3.3 Hz, 1H), 6.58 (d, $J$ = 3.3 Hz, 1H), 4.48 – 4.15 (m, $J$ = 71.8, 67.9, 33.9 Hz, 3H), 2.92 (s, 2H), 2.14 – 2.01 (m, 2H), 1.90 (qd, $J$ = 12.4, 4.2 Hz, 2H), 1.50 (s, 9H).

tert-butyl 4-[5-chloro-4-(6-oxo-2,5-diazaspiro[3.4]octan-2-yl)indol-1-yl]piperidine-1-carboxylate

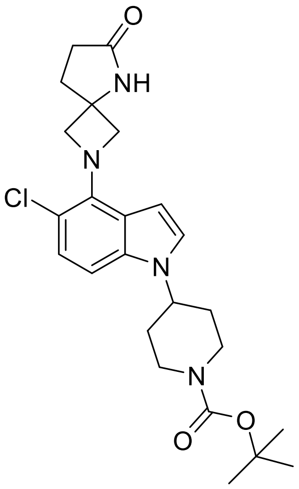

tert-butyl 4-(4-bromo-5-chloro-indol-1-yl)piperidine-1-carboxylate (300 mg, 0.723 mmol), 5-aza-2-azoniaspiro[3.4]octan-6-one;2,2,2-trifluoroacetate (208 mg, 0.868 mmol), RuPhos Pd G3 (60 mg, 0.0723 mmol) and $Cs_2CO_3$ (471 mg, 1.45 mmol) were dissolved in anhydrous dioxane (10mL) under an $N_2$ atmosphere. The reaction mixture was stirred at 100 °C for 18 h. The solvent was removed from the reaction mixture under reduced pressure and the crude material was purified by flash column chromatography (0-20% MeOH in $CH_2Cl_2$) to afford the product tert-butyl 4-[5-chloro-4-(6-oxo-2,5-diazaspiro[3.4]octan-2-yl)indol-1-yl]piperidine-1-carboxylate (65 mg, 17.6%) as a colourless solid.

LCMS: m/z calcd. for [$M+H^+$] 459.2, found: 459.0; RT = 1.81 min.

$^1$H NMR (500 MHz, $CDCl_3$) δ ppm: 7.06 – 7.00 (m, 2H), 6.75 (d, $J$ = 8.7 Hz, 1H), 6.55 (d, $J$ = 3.3 Hz, 1H), 4.58 (d, $J$ = 8.6 Hz, 2H), 4.46 (d, $J$ = 8.6 Hz, 2H), 4.39 – 4.18 (m, 3H), 2.95 – 2.82 (m, 2H), 2.51 – 2.38 (m, 4H), 2.04 (t, $J$ = 6.0 Hz, 2H), 1.84 (app qd, $J$ = 12.4, 4.2 Hz, 2H), 1.49 (s, 9H).

2-(5-chloro-1-(piperidin-4-yl)-1H-indol-4-yl)-2,5-diazaspiro[3.4]octan-6-one (compound **8**)

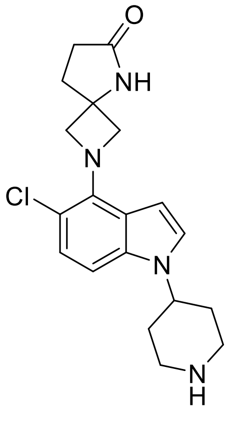

To a solution of tert-butyl 4-[5-chloro-4-(6-oxo-2,5-diazaspiro[3.4]octan-2-yl)indol-1-yl]piperidine-1-carboxylate (1.00 eq, 23 mg, 0.0401 mmol) in DCM (1mL) was added 2,2,2-trifluoroacetic acid (10.0 eq, 0.031 mL, 0.401 mmol). The solution was stirred at rt for 3h. After completeness of the reaction, the solvent was removed under reduced pressure. The crude was purified by preparative HPLC (Waters XSelect CSH C18 ODB column, 5μ silica, 30 mm diameter, 100 mm length), using decreasingly polar mixtures of water (containing 1% formic acid) and MeCN as eluents (gradient 5-95%). Fractions containing the desired compound were evaporated to dryness to afford 2-(5-chloro-1-(piperidin-4-yl)-1H-indol-4-yl)-2,5-diazaspiro[3.4]octan-6-one (4 mg, 20%) as a white solid.

LCMS: m/z calcd. for [M+H$^+$] 359.2, found: 359.1; RT = 1.06 min.

$^1$H NMR (500 MHz, MeOD) δ ppm: 8.55 (s, 1H, NH), 7.22 (d, *J* = 3.5 Hz, 1H), 7.00 (d, *J* = 8.7 Hz, 1H), 6.93 (d, *J* = 8.7 Hz, 1H), 6.68 (d, *J* = 3.5 Hz, 1H), 4.60 – 4.56 (m, 2H), 4.52 (tt, *J* = 11.8, 4.0 Hz, 1H), 4.47 – 4.41 (m, 2H), 3.47 – 3.40 (m, 2H), 3.11 (td, *J* = 13.0, 2.9 Hz, 2H), 2.55 – 2.48 (m, 2H), 2.47 – 2.37 (m, 2H), 2.20 – 2.14 (m, 2H), 2.13 – 2.00 (m, 2H).

7-(1-cyclohexylbenzimidazol-2-yl)-1-methyl-pyrido[2,3-b][1,4]oxazin-2-one (compound S1)

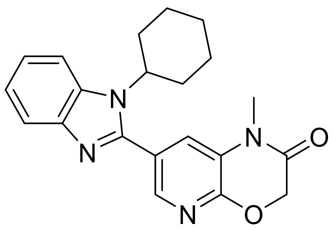

1-Methyl-2-oxo-pyrido[2,3-b][1,4]oxazine-7-carboxylic acid (19 mg, 0.0913 mmol), N2-cyclohexylbenzene-1,2-diamine (22.58 mg, 0.119 mmol), HATU (41.65 mg, 0.11 mmol), and triethylamine (50.6 µL, 0.365 mmol) were dissolved in dimethylformamide (0.5 mL) at room temperature. The reaction mixture was stirred for 5 minutes, followed by the addition of acetic acid (1 mL), and further stirred for 17 hours at room temperature. The reaction progress was monitored by HPLC-MS, confirming complete conversion to the desired product. The reaction mixture was diluted with methanol and water, filtered, and purified by semipreparative HPLC. Suitable fractions were combined and evaporated under reduced pressure which afforded 7-(1-cyclohexylbenzimidazol-2-yl)-1-methyl-pyrido[2,3-b][1,4]oxazin-2-one (12.3 mg, 37% yield).

$^1$H NMR (400 MHz, DMSO-d6) δ ppm: 8.23 (d, *J* = 2.0 Hz, 1H), 8.12-8.18 (m, 1H), 7.81-7.87 (m, 2H), 7.48-7.54 (m, 2H), 5.04 (s, 2H), 4.38 (tt, *J* = 12.3, 3.8 Hz, 2H), 2.19-2.44 (m, 2H), 2.05 (d, *J* = 10.6 Hz, 2H), 1.86 (d, *J* = 11.7 Hz, 2H), 1.61-1.68 (m, 1H), 1.30-1.47 (m, 3H).

7-(1-isopropyl-1H-benzo[d]imidazol-2-yl)-1H-pyrido[2,3-b][1,4]oxazin-2(3H)-one (compound 13)

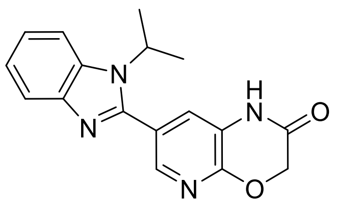

2-oxo-1H,2H,3H-pyrido[2,3-b][1,4]oxazine-7-carboxylic acid (19.8 mg, 0.1 mmol) and N,N-diisopropylethylamine (DIPEA, 51.6 µL, 0.3 mmol) were dissolved in dimethylformamide (2 mL) at room temperature. The mixture was stirred briefly, followed by the addition of HATU (41.8 mg, 0.11 mmol) and N1-(propan-2-yl)benzene-1,2-diamine (15.0 mg, 0.1 mmol). The reaction was stirred for 1 hour at room temperature. LC-MS analysis confirmed the formation of the intermediate product. The reaction mixture was filtered through basic alumina, washed with DMF/MeOH (9:1), and the solvent was removed under reduced pressure. To achieve cyclization, the residue was treated with acetic acid (2 mL) and heated under reflux conditions for 16 hours. LC-MS analysis confirmed the formation of the desired product. The reaction mixture was concentrated, dissolved in DMF, and purified using preparative LC-MS. The purified fractions were lyophilized to afford the product 1H,2H,3H-pyrido[2,3-b][1,4]oxazin-2-one (25.1 mg, 81.4 mmol) as a yellow resin.

$^1$H NMR (400 MHz, DMSO-d6) δ ppm: 11.17 (s, 1H), 8.17 (d, *J* = 2.0 Hz, 1H), 8.03-8.11 (m, 1H), 7.75-7.88 (m, 1H), 7.58 (d, *J* = 2.2 Hz, 1H), 7.48 (dd, *J* = 6.1, 3.2 Hz, 2H), 4.94 (s, 2H), 4.78 (spt, *J* = 13.8 Hz, 1H), 1.66 (d, *J* = 6.8 Hz, 6H).

### Data Tables

#### Table S1. SPR Data Table CRBN$^{midi}$

Affinity of CRBN binders measured by SPR C-avi CRBN$^{midi}$-on-chip, using a steady state affinity model.

| Compound | Compound Class | Kd (nM) | | | | | Assay top concentration (nM) |
|---|---|---|---|---|---|---|---|
| | | 1 | 2 | 3 | Mean | SD | |
| Lenalidomide | IMiD | 135 | 371 | 379 | 295.0 | 138.6 | 500 |
| Compound **9** | DHU | 97.3 | 107 | 119 | 107.8 | 10.9 | 500 |
| Compound **10** | Cyclimid | 32100 | 33800 | 32300 | 32733 | 929.16 | 100000 |
| Compound **11** | Phenylglutarimide | 2560 | 2470 | 2100 | 2376.7 | 243.8 | 10000 |
| Compound **12** | Aminoglutarimide | 2530 | 2510 | 2520 | 2520.0 | 10 | 100000 |
| Compound **8** | 5,4-Spiro | 187 | 194 | 183 | 188.0 | 5.57 | 1000 |
| Compound **4** | Morpholinone | 280 | 270 | 244 | 264.3 | 19.1 | 1000 |
| Compound **S1** | Morpholinone, negative control | 18100 | 19200 | 19300 | 18900 | 665.8 | 100000 |

#### Table S2. ITC measurements of CRBN binders with CRBN:DDB1$^{\Delta BPB}$

| Compound | Compound Class | Measured Parameters | | | | Classification |
|---|---|---|---|---|---|---|
| | | $K_D$ (μM) | N | ΔH (kJ/mol) | −TΔS (kJ/mol) | |
| Lenalidomide | IMiD | 0.223 ± 0.025 | 1.78 | 10.8 ± 0.074 | −48.8 | exothermic |
| **4** | Morpholinone | 0.899 ± 0.055 | 1.66 | 29.8 ± 0.186 | −4.74 | endothermic |
| **8** | 5,4-Spiro | 2.02 ± 0.102 | 1.24 | −28.8 ± 0.238 | −3.74 | endothermic |
| **9** | DHU | 0.039 ± 0.007 | 1.08 | 9.77 ± 0.123 | −52.1 | exothermic |
| **11** | Phenylglutarimide | 8.51 ± 0.59 | 1.42 | 19.3 ± 0.405 | −48.3 | exothermic |

Table S3. Measurement of different CRBN classes by DSF with CRBN$^{midi}$. Turbidity first derivative of thermal denaturation for CRBN$^{midi}$ in the absence or presence of binders.

| Compound | Compound Class | CRBN$^{midi}$ Tm, turbidity (°C) | | | | | | CRBN$^{TBD}$ |
|---|---|---|---|---|---|---|---|---|
| | | 1 | 2 | 3 | Mean | SD | Average Δ Tm | Average Δ Tm |
| Apo | / | 44.07 | 44.13 | 44.05 | 44.08 | 0.04 | / | / |
| Lenalidomide | IMiD | 58.03 | 58.04 | 58.11 | 58.05 | 0.05 | 13.97 | 3.6 |
| **4** | Morpholinone | 41.15 | 41.16 | 41.21 | 41.17 | 0.03 | -2.91 | 7.4 |
| **8** | 5,4-Spiro | 47.18 | 47.38 | 47.29 | 47.29 | 0.10 | 3.21 | 6.1 |
| **9** | Dihydrouracil | 58.61 | 58.65 | 58.67 | 58.65 | 0.03 | 14.57 | 5.1 |
| **10** | Cyclimid | 55.00 | 54.92 | 54.95 | 54.96 | 0.04 | 10.88 | 0.6 |
| **11** | Phenylglutarimide | 55.96 | 56.14 | 56.01 | 56.03 | 0.09 | 11.95 | 2.6 |
| **12** | Aminoglutarimide | 55.47 | 55.58 | 55.41 | 55.49 | 0.08 | 11.41 | n.d. |

Table S4. Turbidity first derivative of thermal denaturation for CRBN$^{midi}$ WT and mutants in the absence or presence of binders. WT is mean of triplicates.

| Mutation | Compound | Compound class | Tm | ΔTm |
|---|---|---|---|---|
| WT | Apo | / | 44.08 | - |
| WT | Lenalidomide | Imid | 58.05 | 13.97 |
| WT | Compound 9 | DHU | 58.65 | 14.57 |
| WT | Compound 8 | 5,4-Spiro | 47.29 | 3.21 |
| WT | Compound 4 | Morpholinone | 41.17 | -2.91 |
| H378N | Apo | / | 43.00 | - |
| H378N | Lenalidomide | Imid | 55.49 | 12.49 |
| H378N | Compound 9 | DHU | 56.14 | 13.14 |
| H378N | Compound 8 | 5,4-Spiro | 44.70 | 1.70 |
| H378N | Compound 4 | Morpholinone | 41.65 | -1.35 |
| H378A | Apo | / | 43.05 | - |
| H378A | Lenalidomide | Imid | 53.99 | 10.94 |
| H378A | Compound 9 | DHU | 53.91 | 10.86 |
| H378A | Compound 8 | 5,4-Spiro | 47.65 | 4.6 |
| H378A | Compound 4 | Morpholinone | 41.81 | -1.24 |

| Mutation | Compound | Compound class | Tm | ΔTm |
|---|---|---|---|---|
| Q100A | Apo | / | 44.00 | - |
| Q100A | Lenalidomide | Imid | 58.00 | 14.00 |
| Q100A | Compound 9 | DHU | 58.75 | 14.75 |
| Q100A | Compound 8 | 5,4-Spiro | 47.17 | 3.17 |
| Q100A | Compound 4 | Morpholinone | 41.37 | -2.63 |
| L60A | Apo | / | 44.31 | - |
| L60A | Lenalidomide | Imid | 51.38 | 7.07 |
| L60A | Compound 9 | DHU | 49.53 | 5.22 |
| L60A | Compound 8 | 5,4-Spiro | 46.47 | 2.16 |
| L60A | Compound 4 | Morpholinone | 41.62 | -2.69 |
| L60A H378A | Apo | / | 42.80 | - |
| L60A H378A | Lenalidomide | Imid | 49.23 | 6.43 |
| L60A H378A | Compound 9 | DHU | 47.45 | 4.65 |
| L60A H378A | Compound 8 | 5,4-Spiro | 43.79 | 0.99 |
| L60A H378A | Compound 4 | Morpholinone | 39.83 | -2.97 |

#### Table S5. Small Angle X-ray scattering statistics for $CRBN^{midi}$ bound to each binder series

| SASBDB ID | X | X | X | X | X | X |
|---|---|---|---|---|---|---|
| Sample Details | | | | | | |
| Organism | *E. coli* BL21 (DE3) | | | | | |
| Source | Recombinantly expressed | | | | | |
| Uniprot sequence ID | *Derived from human Cereblon, Uniprot Q96SW2* | | | | | |
| Description | $CRBN^{midi}$ | $CRBN^{midi}$ bound to lenalidomide | $CRBN^{midi}$ bound to Compound **9** | $CRBN^{midi}$ bound to Compound **11** | $CRBN^{midi}$ bound to Compound **8** | $CRBN^{midi}$ bound to Compound **4** |
| Expected molecular mass (kDa) | 37.4 | 37.7 | 37.7 | 37.7 | 37.7 | 37.7 |
| Loading concentration (mg/mL) | 3.4 | | | | | |
| Injection volume (μL) | 50 | | | | | |
| Buffer composition | 20 mM HEPES pH 7.5, 500 mM NaCl, 0.5 mM TCEP | | | | | |
| SAS data collection parameters | | | | | | |
| Source and instrument | B21, Diamond Light Source | | | | | |
| Wavelength (Å) | 0.9464 | | | | | |
| Sample-detector distance (m) | 3.7 | | | | | |
| q-measurement range (Å) | $4.5 \times 10^{-3}$- $3.4 \times 10^{-1}$ | | | | | |
| Radiation damage monitoring | Frame-by-frame comparison | | | | | |
| Exposure time (s) and number | 3 x 600 | | | | | |
| Sample configuration | SEC-SAXS, Cytiva Superdex 200 Increase 3.2/300 | | | | | |
| Sample temperature (°C) | 15 | | | | | |
| Structural parameters | | | | | | |
| Guinier analysis | | | | | | |
| I(0) ($cm^{-1}$) | $0.017 \pm 3.4 \times 10^{-5}$ | $0.017 \pm 2.4 \times 10^{-5}$ | $0.017 \pm 2.8 \times 10^{-5}$ | $0.012 \pm 1.6 \times 10^{-5}$ | $0.021 \pm 2.6 \times 10^{-5}$ | $0.0096 \pm 2.4 \times 10^{-5}$ |
| $R_g$ (Å) | 26.83 ± 0.09 | 22.99 ± 0.06 | 23.43 ± 0.07 | 22.80 ± 0.06 | 26.04 ± 0.06 | 25.72 ± 0.11 |
| $qR_g$ max | 1.28 | 1.30 | 1.30 | 1.30 | 1.30 | 1.30 |
| P(r) analysis from AUTOGNOM | | | | | | |
| I(0) ($cm^{-1}$) | $0.016 \pm 2.7 \times 10^{-5}$ | $0.017 \pm 1.9 \times 10^{-5}$ | $0.017 \pm 2.8 \times 10^{-5}$ | $0.012 \pm 1.5 \times 10^{-5}$ | $0.021 \pm 2.5 \times 10^{-5}$ | $0.0095 \pm 2.2 \times 10^{-5}$ |
| $R_g$ (Å) | 26.26 ± 0.051 | 22.56 ± 0.035 | 23.11 ± 0.054 | 22.55 ± 0.046 | 26.05 ± 0.0049 | 25.43 ± 0.060 |
| $d_{max}$ (Å) | 154.26 | 137.92 | 141.94 | 141.42 | 161.52 | 145.16 |
| $q$ range ($Å^{-1}$) | 0.0126-0.2982 | 0.0089-0.3400 | 0.0093-0.3400 | 0.0111-0.3400 | 0.0101-0.3071 | 0.0126-0.3110 |
| $\chi^2$ (total estimate from *GNOM*) | 0.87 | 0.86 | 0.81 | 0.78 | 0.83 | 0.85 |

| | | | | | | |
|---|---|---|---|---|---|---|
| Porod volume (Å$^{-3}$) (ratio $V_p$/calculated $M$) | 70306.90 | 59262.70 | 61505.60 | 63949 | 60447.10 | 55533.10 |

#### Table S6. Small Angle X-ray Scattering statistics for CRBN$^{midi}$ mutants bound to compound 9

| SASBDB ID | X | X | X | X | X |
|---|---|---|---|---|---|
| **Sample Details** | | | | | |
| Organism | *E. coli* BL21 (DE3) | | | | |
| Source | Recombinantly expressed | | | | |
| Uniprot sequence ID | *Derived from human Cereblon, Uniprot Q96SW2* | | | | |
| Description | CRBN$^{midi}$ H378A | CRBN$^{midi}$ H378N | CRBN$^{midi}$ L60A | CRBN$^{midi}$ L60A H378A | CRBN$^{midi}$ Q100A |
| Expected molecular mass (kDa) | 37.4 | 37.7 | 37.7 | 37.6 | 37.7 |
| Loading concentration (mg/mL) | 3.4 | | | | |
| Injection volume (µL) | 50 | | | | |
| Buffer composition | 20 mM HEPES pH 7.5, 500 mM NaCl, 0.5 mM TCEP | | | | |
| **SAS data collection parameters** | | | | | |
| Source and instrument | B21, Diamond Light Source | | | | |
| Wavelength (Å) | 0.9464 | | | | |
| Sample- detector distance (m) | 3.7 | | | | |
| q-measurement range (Å) | $4.5 \times 10^{-3}$- $3.4 \times 10^{-1}$ | | | | |
| Radiation damage monitoring | Frame-by-frame comparison | | | | |
| Exposure time (s) and number | 3 x 600 | | | | |
| Sample configuration | SEC-SAXS, Cytiva Superdex 200 Increase 3.2/300 | | | | |
| Sample temperature (°C) | 15 | | | | |
| **Structural parameters** | | | | | |
| **Guinier analysis** | | | | | |
| I(0) ($cm^{-1}$) | $0.012 \pm 1.6 \times 10^{-5}$ | $0.013 \pm 2.9 \times 10^{-5}$ | $0.012 \pm 1.8 \times 10^{-5}$ | $0.014 \pm 2.0 \times 10^{-5}$ | $0.011 \pm 1.7 \times 10^{-5}$ |
| $R_g$ (Å) | 22.88 ± 0.06 | 23.80 ± 0.10 | 24.84 ± 0.07 | 25.18 ± 0.06 | 23.07 ± 0.07 |
| $qR_g$ max | 1.30 | 1.30 | 1.29 | 1.30 | 1.30 |
| **P(r) analysis from AUTOGNOM** | | | | | |
| I(0) ($cm^{-1}$) | $0.011 \pm 1.5 \times 10^{-5}$ | $0.012 \pm 2.3 \times 10^{-5}$ | $0.012 \pm 1.8 \times 10^{-5}$ | $0.014 \pm 1.7 \times 10^{-5}$ | $0.011 \pm 1.3 \times 10^{-5}$ |
| $R_g$ (Å) | 22.35 ± 0.033 | 22.83 ± 0.041 | 24.56 ± 0.054 | 25.05 ± 0.039 | 22.38 ± 0.032 |
| $d_{max}$ (Å) | 124.76 | 123.82 | 145.48 | 149.88 | 125.78 |
| $q$ range (Å$^{-1}$) | 0.0083-0.3400 | 0.0091-0.3361 | 0.0121-0.3220 | 0.0090-0.3177 | 0.0082-0.3400 |
| $\chi^2$ (total estimate from *GNOM*) | 0.87 | 0.85 | 0.82 | 0.86 | 0.86 |
| Porod volume (Å$^{-3}$) (ratio $V_p$/calculated $M$) | 54576 | 64098 | 61883 | 63723 | 56333 |

#### Table S7. X-Ray Crystallographic Data and Refinement Statistics Data

| | $CRBN^{midi}$:Compound **9** |
|---|---|
| **Data collection** | |
| Space group | $P12_11$ |
| Cell dimensions | |
| *a*, *b*, *c* (Å) | 51.42, 153.11, 54.05 |
| α, β, γ (°) | 90.00, 118.40, 90.00 |
| Resolution (Å) | 47.54-2.36 (2.56-2.36) |
| $R_{merge}$ | 0.159 (1.459) |
| *I* / s*I* | 7 (1.6) |
| $CC_{1/2}$ | 0.994 (0.657) |
| Completeness (%) (spherical) | 59.2 (13.6) |
| Completeness (%) (ellipsoidal) | 87.1 (70.7) |
| Redundancy | 6.3 (5.8) |
| **Refinement** | |
| Resolution (Å) | 47.54-2.36 |
| No. reflections | 17700 (179) |
| $R_{work}$ / $R_{free}$ | 0.233/ 0.296 |
| No. atoms | 5041 |
| Protein | 4978 |
| Ligand/ion | 48 |
| Water | 15 |
| *B*-factors | |
| Protein | 47.11 |
| Ligand/ion | 37.74 |
| Water | 35.31 |
| R.m.s. deviations | |
| Bond lengths (Å) | 0.018 |
| Bond angles (°) | 0.49 |
| **PDB ID** | 9SUN |

#### Table S8. Cryo-EM data collection, processing and refinement statistics for dihydrouracil **9**

| | Cryo-EM structure of CRBN bound to 1-[1-(4-piperidyl)indol-4-yl]hexahydropyrimidine-2,4-dione in the closed conformation | | | Cryo-EM structure of CRBN bound to yl]hexahydropyrimidine-2,4-dione in th | |
|---|---|---|---|---|---|
| | Composite map<br>EMD-55258<br>PDB 9SVH | Consensus map<br>EMD-55057 | Focused map<br>EMD-55058 | Composite map<br>EMD-55259<br>PDB 9SVI | Consensus map<br>EMD-55059 |
| **Data collection and processing** | | | | | |
| Magnification | 190,000 | | | | |
| Voltage (kV) | 200 | | | | |
| Electron exposure (e–/Å$^2$) | 21.94 | | | | |
| Defocus range (μm) | -(1.7-3.2) | | | | |
| Pixel size (Å) | 0.71 | | | | |
| Symmetry imposed | C1 | | | | |
| Initial particle images (no.) | 1,585,848 | | | | |
| Final particle images (no.) | 56,510 | | | 86,485 | |
| Map resolution (Å) | | 3.66 | 4.07 | | 3.33 |
| FSC threshold | | 0.143 | 0.143 | | 0.143 |
| **Refinement** | | | | | |
| Map sharpening *B* factor (Å$^2$) | | 144.2 | 209.9 | | 141.6 |
| Model composition | | | | | |
| Non-hydrogen atoms | 9,164 | | | 8,846 | |
| Protein residues | 1,174 | | | 1,183 | |
| Ligands | A1JP8, ZN | | | A1JP8, ZN | |
| Model-Map | | | | | |
| CC (mask) | 0.69 | | | 0.76 | |
| R.m.s. deviations | | | | | |
| Bond lengths (Å) | 0.004 | | | 0.003 | |
| Bond angles (°) | 0.630 | | | 0.622 | |
| Validation | | | | | |
| MolProbity score | 1.65 | | | 1.45 | |
| Clashscore | 6.29 | | | 4.20 | |
| CaBLAM outliers (%) | 1.48 | | | 1.89 | |
| Rotamer outliers (%) | 0.00 | | | 0.00 | |
| Cβ outliers (%) | 0.00 | | | 0.00 | |
| Ramachandran plot | | | | | |
| Favored (%) | 95.69 | | | 96.26 | |
| Allowed (%) | 4.31 | | | 3.66 | |
| Disallowed (%) | 0.00 | | | 0.09 | |

#### Table S9. Cryo-EM data collection, processing and refinement statistics for spirolactam **8**

| | Cryo-EM structure of CRBN bound to 2-[5-chloro-1-(4-piperidyl)indol-4-yl]-2,5-diazaspiro[3.4]octan-6-one in the open conformation | | |
|---|---|---|---|
| | Composite map<br>EMD-55257<br>PDB 9SVG | Consensus map<br>EMD-54624 | Focused map<br>EMD-54644 |
| **Data collection and processing** | | | |
| Magnification | | 165,000 | |
| Voltage (kV) | | 300 | |
| Electron exposure (e–/Å$^2$) | | 50 | |
| Defocus range (μm) | | -(0.8-1.8) | |
| Pixel size (Å) | | 0.75 | |
| Symmetry imposed | | C1 | |
| Initial particle images (no.) | | 2,575,103 | |
| Final particle images (no.) | | 573,499 | |
| Map resolution (Å) | | 2.22 | 3.44 |
| FSC threshold | | 0.143 | 0.143 |
| **Refinement** | | | |
| Map sharpening *B* factor (Å$^2$) | | 69 | 145.1 |
| Model composition | | | |
| Non-hydrogen atoms | 8,880 | | |
| Protein residues | 1158 | | |
| Ligands | A1JQZ, ZN | | |
| Model-Map | | | |
| CC (mask) | 0.67 | | |
| R.m.s. deviations | | | |
| Bond lengths (Å) | 0.003 | | |
| Bond angles (°) | 0.681 | | |
| Validation | | | |
| MolProbity score | 1.46 | | |
| Clashscore | 4.78 | | |
| CaBLAM outliers (%) | 2.12 | | |
| Rotamer outliers (%) | 0.11 | | |
| Cβ outliers (%) | 0.00 | | |
| Ramachandran plot | | | |
| Favored (%) | 96.68 | | |
| Allowed (%) | 3.23 | | |
| Disallowed (%) | 0.09 | | |

#### Table S10. List of primers for sire-directed mutagenesis

| Primer Name | Sequence 5’-3’ | Mutation introduced |
|---|---|---|
| L60A | CCGACATCACACACCTACGCGGGTGCCGATATGGAAG | Mutate Leu60 to Ala in CRBN$^{midi}$ |
| Q100A | CAGACCTTACCGCTGGCGCTGTTTCACCCGCAGGAAG | Mutate Gln100 to Ala in CRBN$^{midi}$ |
| H378A | CGCCCGTCTACCGAAGCCAGCTGGTTTCCAGGGTATG | Mutate His378 to Ala in CRBN$^{midi}$ and CRBN$^{midiL60A}$ |
| H378N | CGCCCGTCTACCGAAAACAGCTGGTTTCCAGGGTATG | Mutate His378 to Asn in CRBN$^{midi}$ |

#### Table S11. List of cycling parameters for site-directed mutagenesis

| Segment | Cycles | Temperature (°C) | Time (minutes) |
|---|---|---|---|
| 1 | 1 | 95 | 1 |
| 2 | 15 | 95 | 0.5 |
| | | 55 | 1 |
| | | 68 | 8.5 |

#### Table S12. Intact protein LC-MS of CRBN$^{midi}$ mutants

| Mutant | Predicted Mass (Da) | Experimental Mass (Da) |
|---|---|---|
| WT | 37435.88 | 37428.77 |
| H378N | 37412.84 | 37406.20 |
| H378A | 37369.82 | 37362.47 |
| Q100A | 37378.83 | 37370.89 |
| L60A | 37393.80 | 37386.22 |
| L60A H378A | 37327.74 | 37318.07 |

#### Table S13. Primers for cloning CRBN:DDB1$^{\Delta BPB}$

| Primer Name | Sequence |
|---|---|
| DDB1delBPBvec_F | 5’GGAAACGGTAACTCTGGAGAGATCCAGAAGCTGCACATCCGTACCGTGCCTC 3’ |
| DDB1delBPBvec_R | 3’GCTTTTGAGCGGTGACCACGTAGTTGTAGGA 5’ |
| DDB1delBPBins_F | 5’TCCTACAACTACGTGGTCACCGCTCAAAAGC 3’ |
| DDB1delBPBins_R | 3’TCCAGAGTTACCGTTTCCACCGATACCGTTACGGATGATACGCAGGG 5’ |

### Supplementary Figures

#### Figure S1. 2D $^{15}N$ HSQC CSP detected with thalidomide and 15N-labelled $CRBN^{TBD}$

Chemical shift perturbation and line broadening induced upon binding of thalidomide to CRBN TBD. Circles indicate cross peaks affected by the FBS hit depicted in Fig. 2 suggesting that both compounds interact with the same binding site

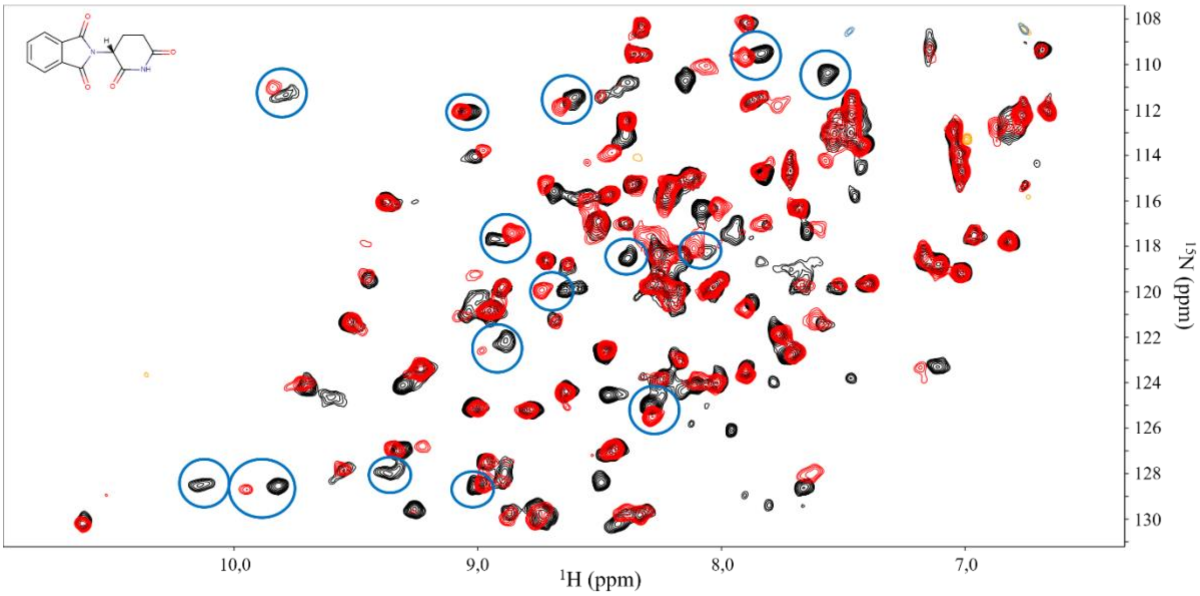

#### Figure S2. $CRBN^{midi}$:Dihydrouracil **9** crystal structure domain swap and comparison to other structures.

Structure of $CRBN^{midi}$ in complex with **9**. $CRBN^{midi}$ is shown in cartoon representation coloured by domain, Lon (yellow), HB (orange), TBD (salmon), GSG linker (cyan), **9** is shown as black sticks. A) The left panel shows the structure with the domain swap modelled and the right panel shows the structure without the domain swap modelled. The GSG linker is shown in stick form and coloured cyan, the polder OMIT (Fo-Fc) map of the linker contoured to 3 σ is shown as green mesh. B) Superposition of TBD (chain A) of $CRBN^{midi}$:**9** with the crystal structure of $CRBN^{TBD}$:compound **C3** (grey, PDB ID: 9ODS). Chemical structures of compound **C3** and DHU **9** are show below.

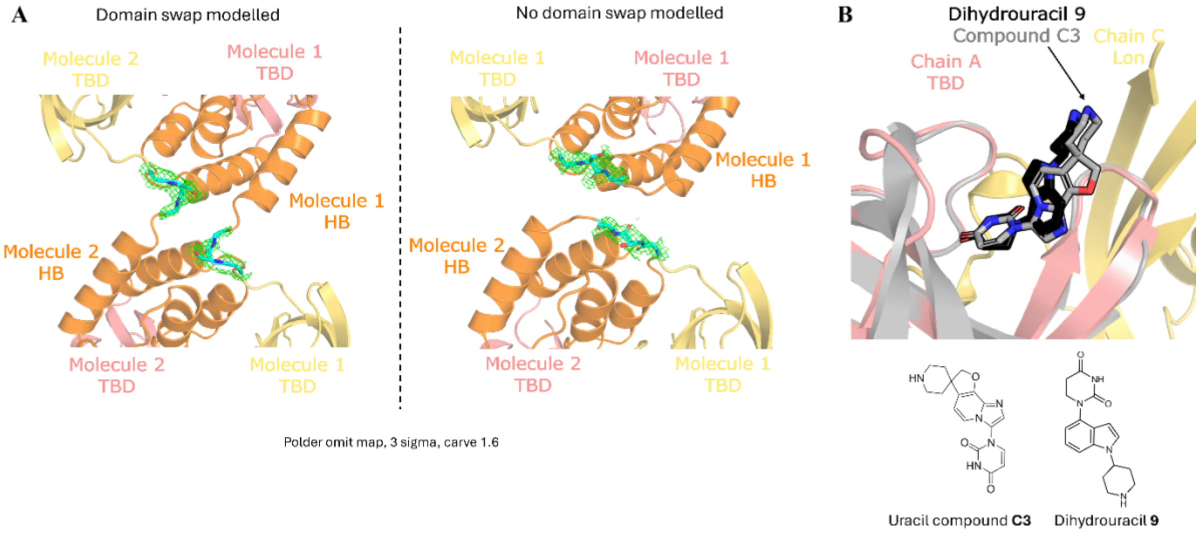

Figure S3. Cryo-EM Workflow for dihydrouracil **9**

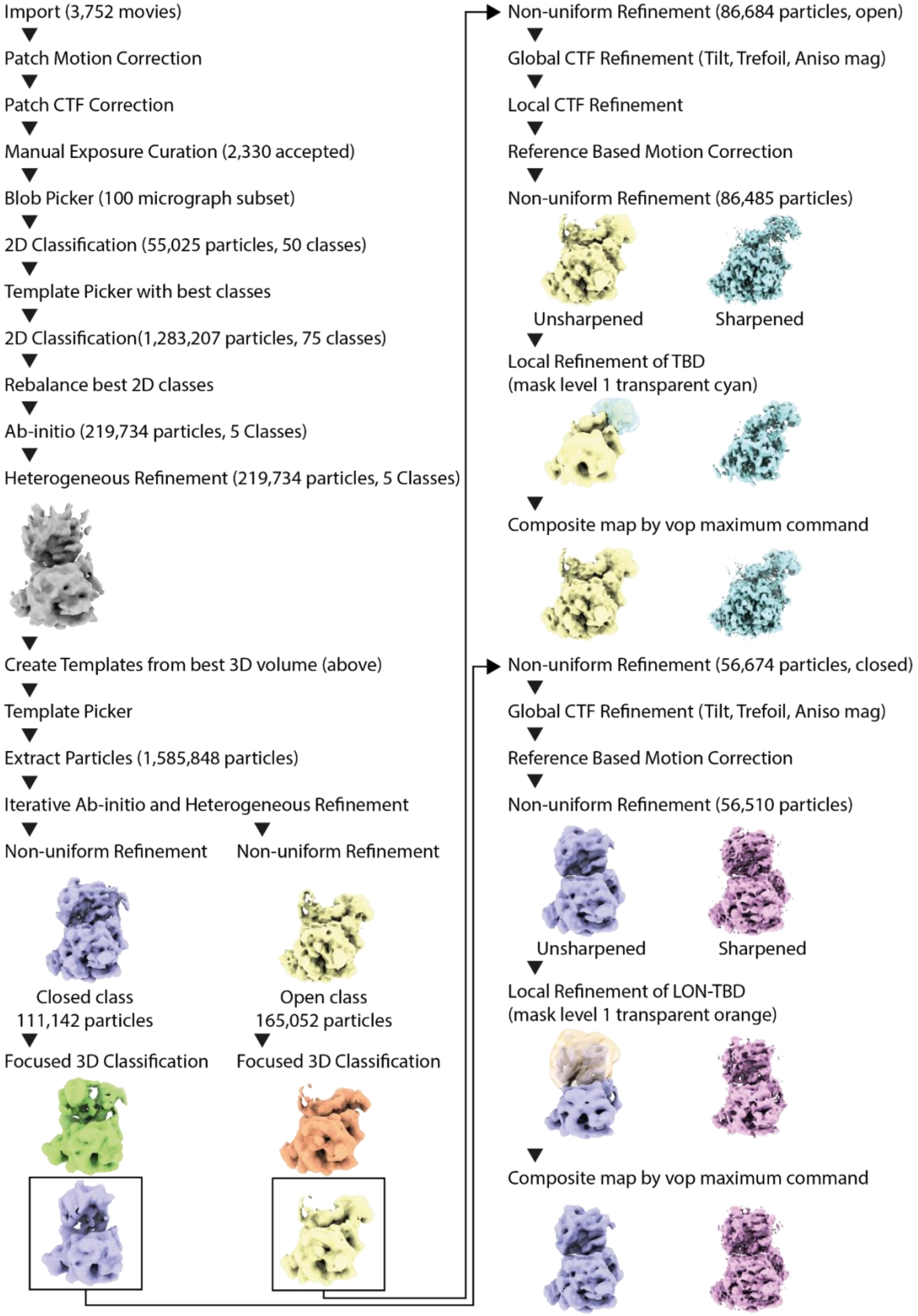

#### Figure S4. Cryo-EM Map Information for dihydrouracil **9**

Workflow for Cryo-EM data processing of dataset for CRBN-DDB1ΔBPB bound to dihydrouracil ligand 1-[1-(4-piperidyl)indol-4-yl]hexahydropyrimidine-2,4-dione. Gold-standard Fourier shell correlation, cFSC, angular distribution, and posterior position directional distribution plots for the consensus and locally refined maps of CRBN-DDB1ΔBPB bound to dihydrouracil ligand 1-[1-(4-piperidyl)indol-4-yl]hexahydropyrimidine-2,4-dione in the open and closed conformation.

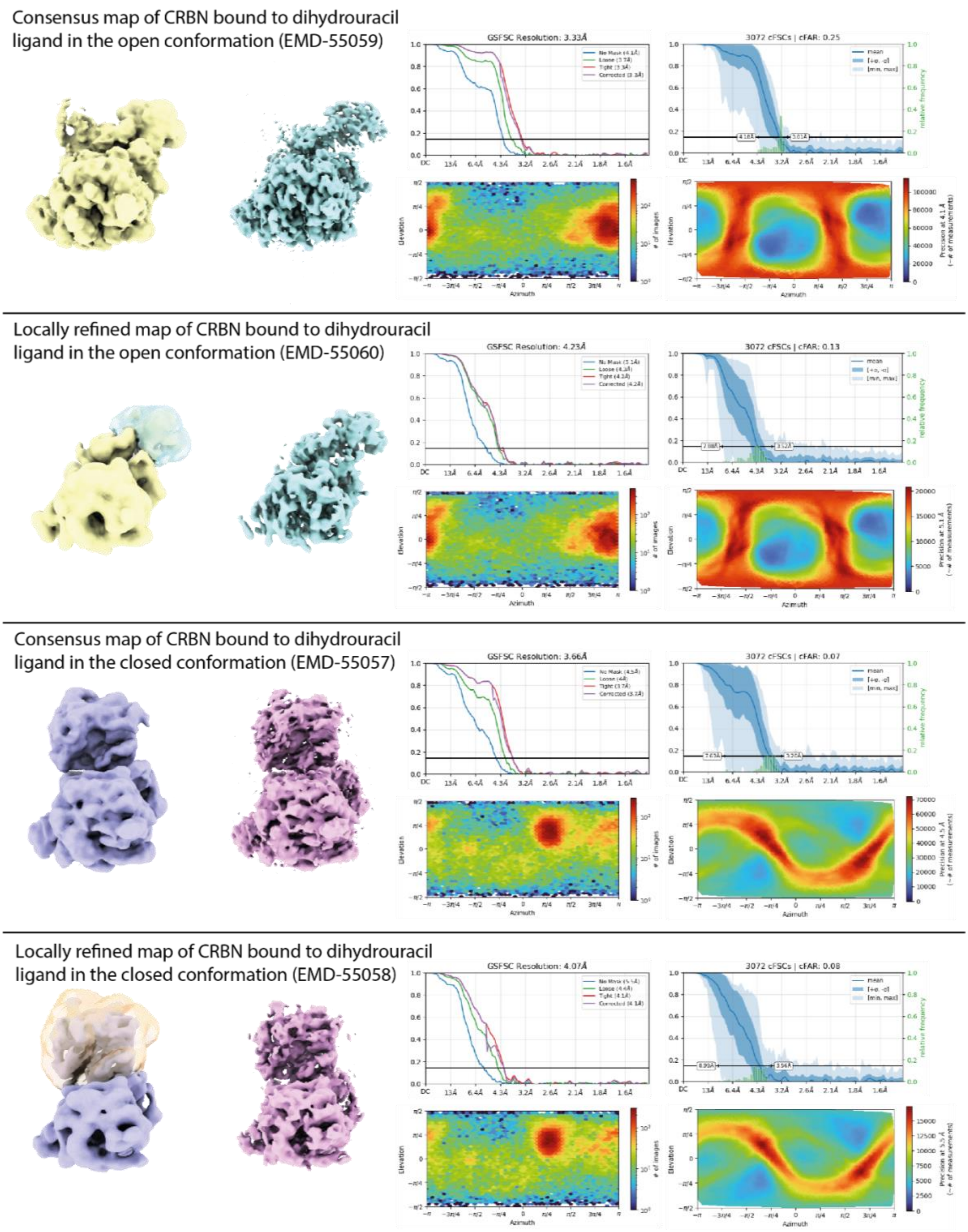

#### Figure S5. Cryo-EM Workflow for 5,4-spiro **8**

Workflow for Cryo-EM data processing of dataset for CRBN-DDB1ΔBPB bound to spirocyclic ligand 2-[5-chloro-1-(4-piperidyl)indol-4-yl]-2,5-diazaspiro[3.4]octan-6-one2-[5-chloro-1-(4-piperidyl)indol-4-yl]-2,5-diazaspiro[3.4]octan-6-one.

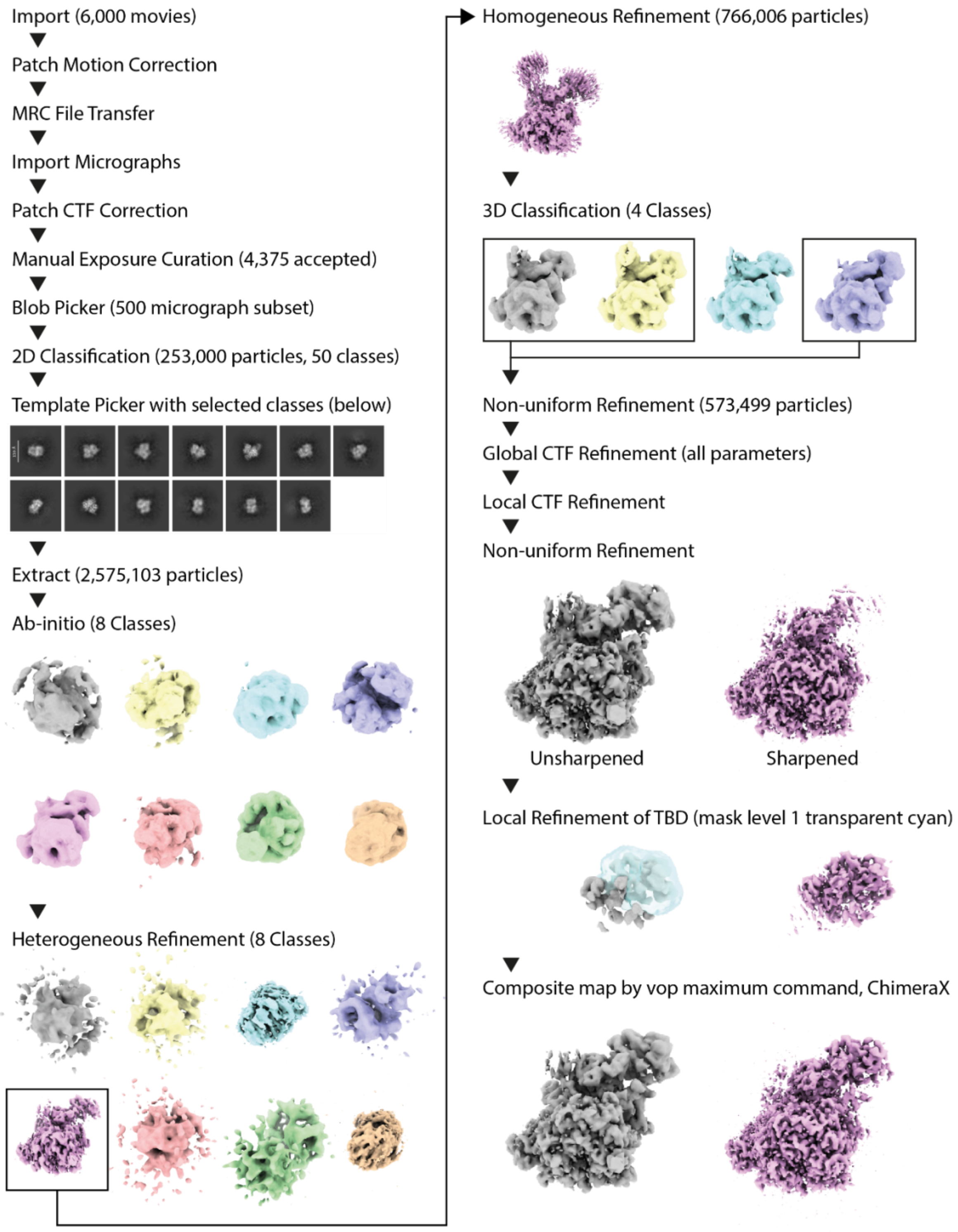

#### Figure S6. Cryo-EM Map Information for 5,4-spiro **8**

Gold-standard Fourier shell correlation, cFSC, angular distribution, and posterior position directional distribution plots for the consensus and locally refined maps of CRBN-DDB1ΔBPB bound to spirocyclic ligand 2-[5-chloro-1-(4-piperidyl)indol-4-yl]-2,5-diazaspiro[3.4]octan-6-one2-[5-chloro-1-(4-piperidyl)indol-4-yl]-2,5-diazaspiro[3.4]octan-6-one.

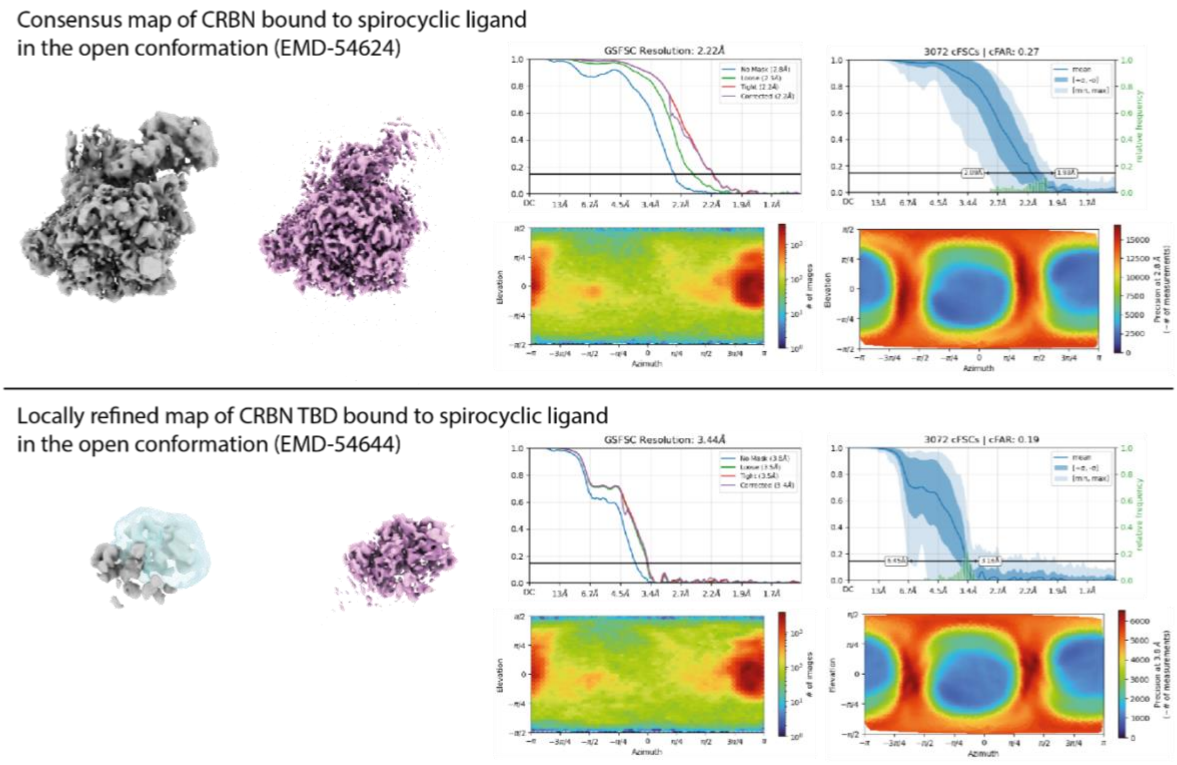

#### Figure S7. 5,4-spiro **8** Binding Site in Locally Refined Cryo-EM Map

Locally refined map of TBD bound to spirolactam 8 showing binding site.

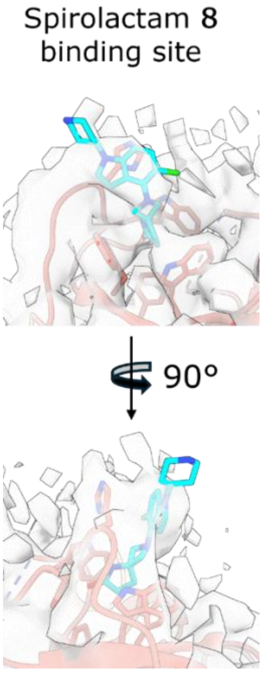

#### Figure S8. Validating the stability of FLAG-CRBN expressed into DLD-1 CRBN KO background cells.

A

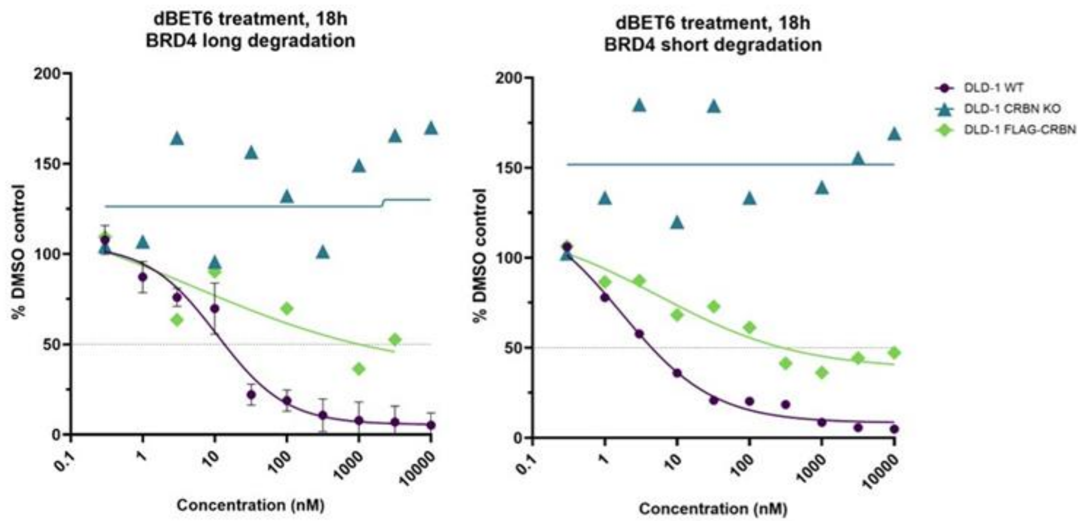

B

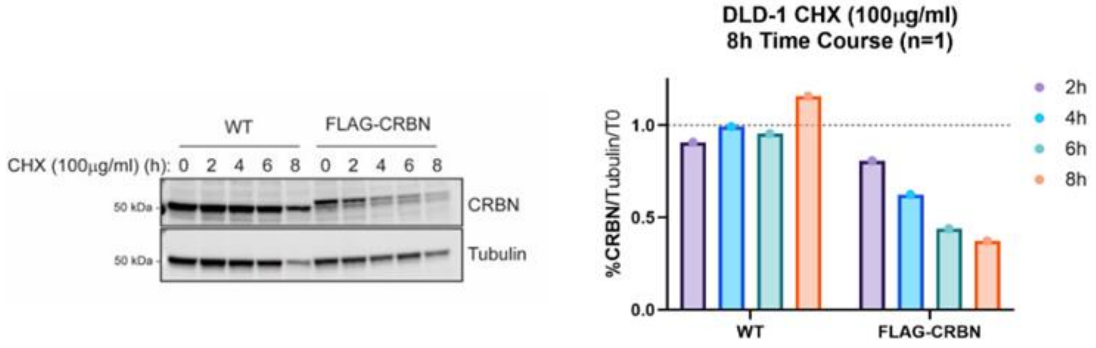

Cells were treated with 100 mg/ml cycloheximide (CHX) over an 8h time-course. CRBN levels were quantified and normalised against loading control and time zero (T0) expression.

#### Figure S9. BRD4 Degradation in DLD-1 WT, CRBN KO and CRBN KO stably expressing FLAG-CRBN upon Treatment with dBET6

Representative western blots showing degradation of BRD4 in (A) DLD-1 wild-type (WT), (B) CRBN-KO and (C) CRBN KO stably expressing FLAG-CRBN. Cells were treated with dBET6 for 18h before analysis by western blotting. Quantification of all replicates is summarised in Figure S8A.

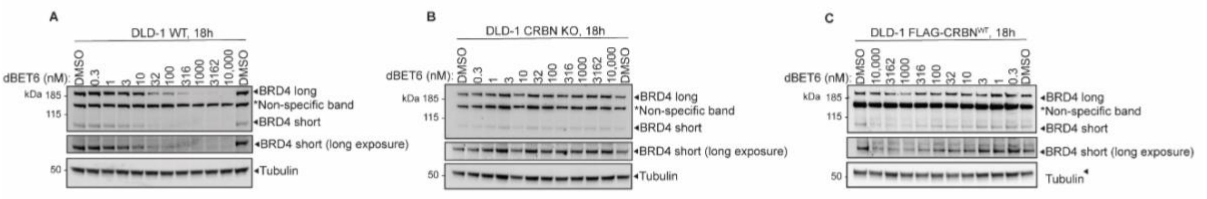
